# *CTGoMartini*: A Python Framework for Simulating Biomolecular Conformational Transitions with Gō-Martini Models

**DOI:** 10.64898/2026.04.30.721921

**Authors:** Song Yang, Chen Song

## Abstract

Characterizing conformational transitions between distinct structural states is essential for understanding protein function but remains challenging due to the timescale limitations of atomistic molecular dynamics. While coarse-grained models like Martini accelerate sampling, classical elastic-network or Gō-like restraints often trap proteins in a single energy basin, precluding the study of transition pathways between distinct functional states. Here, we present CTGoMartini, a comprehensive Python package designed to simulate protein conformational transitions using Gō-Martini models in explicit membranes. CTGoMartini addresses key methodological limitations of existing approaches by redefining native contacts as a dedicated interaction type, thereby eliminating spurious protein aggregation artifacts in multi-copy simulations. The package implements both switching and multiple-basin approaches (Exponential and Hamiltonian mixing) to sample transitions between experimentally defined states. Furthermore, it integrates Hamiltonian replica exchange molecular dynamics (HREMD) with PyMBAR analysis, enabling efficient optimization of mixing parameters that govern barrier heights and relative state stabilities. We demonstrate the power of CTGoMartini through two biologically significant membrane protein systems: (1) capturing the inward-open to outward-open transition of the lipid transporter SPNS2, revealing the molecular mechanism of S1P translocation; and (2) elucidating how membrane surface tension and anionic lipids (POPA, PIP_2_) modulate the conformational equilibrium of the mechanosensitive ion channel TREK1. By streamlining model construction, simulation, and analysis, CTGoMartini offers an easy-to-use platform that connects static structural snapshots with their underlying dynamic functional mechanisms.

**TOC Graphic:** 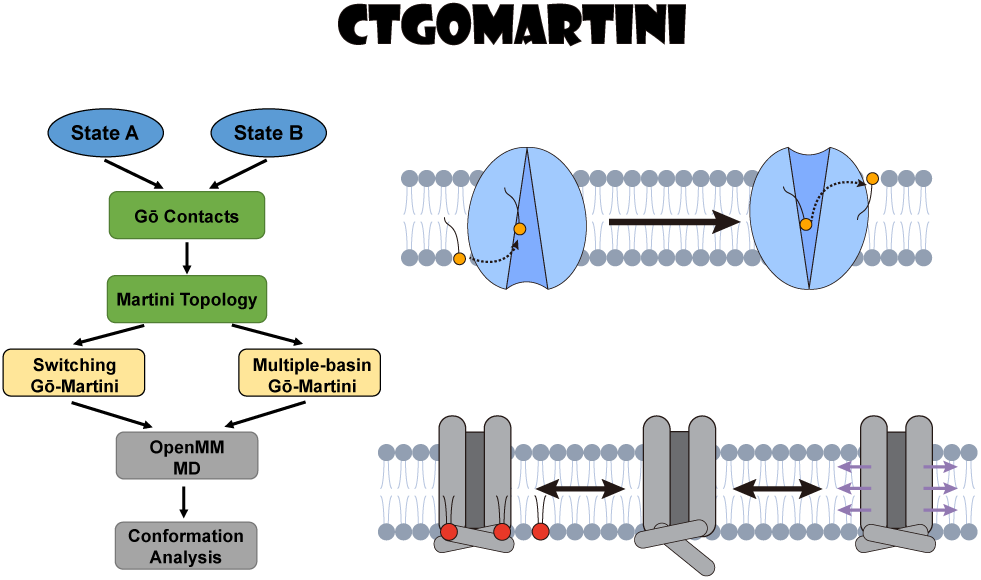

## Introduction

Proteins are dynamic molecular machines that often rely on conformational transitions between distinct structural states to perform their biological functions. These transitions underpin essential cellular processes. For example, membrane transporters, such as the glucose transporter GLUT1, undergo large-scale rearrangements between inward-open and outward-open states to shuttle substrates across the membrane. ^1^ Similarly, G protein-coupled receptors (GPCRs) like the β_2_-adrenergic receptor transition between inactive and active states to transmit extracellular signals into the cell. ^2^ Malfunctions in these precisely orchestrated transitions are linked to numerous diseases, including cystic fibrosis (CFTR channel), ^3^ neurodegenerative disorders (misfolded proteins), ^4^ and cancer (signaling kinases). ^5^ Therefore, characterizing the pathways and energetics of these transitions at atomic detail is critical not only for a fundamental mechanistic understanding of protein functions but also for the rational design of drugs that can modulate specific conformational states.

Experimentally, techniques such as cryo-electron microscopy (cryo-EM) and X-ray crystallography can provide high-resolution structural snapshots of different states. However, they often struggle to capture the continuous, transient pathways that connect these states. ^6^ Deeplearning methods, including AlphaFold2^7^ and AlphaFold3, ^8^ have revolutionized structure prediction, but they are primarily designed to predict thermodynamically stable conformations rather than to model protein dynamics. While emerging approaches such as BioEmu ^9^ are beginning to sample conformational ensembles of proteins, they remain largely unable to capture protein–environment interactions, especially for protein–lipid interactions.

Molecular dynamics (MD) simulation serves as a powerful computational microscope providing high-resolution spatial and temporal insights into biomolecular motion. ^10,11^ However, a fundamental challenge stems from the disparity in timescales. While many functional conformational changes occur on the millisecond regime, all-atom MD simulations with explicit solvent are typically limited to microseconds for systems of biologically relevant size, despite continuous advances in hardware and algorithms. ^12^ This gap hinders the direct observation of many large-scale, functionally important transitions. Coarse-grained (CG) models address this challenge by reducing the number of degrees of freedom, thereby accelerating conformational sampling by several orders of magnitude. The Martini force field, particularly its recent version 3, has emerged as a widely adopted CG model for biomolecular simulations, offering a balanced representation of chemical specificity and computational efficiency. ^13^ In standard Martini protein simulations, an elastic network model (ENM) restraint is applied to the protein backbone to maintain the secondary or higher fold. ^14^ Although effective in stabilizing a single native state, the harmonic potentials of the ENM intrinsically restrict large-scale conformational changes, effectively locking the protein in a single energy basin. To overcome this limitation and enable the study of conformational transitions, structure-based Gō-models have been integrated with the Martini framework. ^15,16^ Gō-models bias the energy landscape towards a set of predefined native contacts derived from experimental structures. By defining multiple sets of native contacts corresponding to different known conformations (e.g., an inward-open and an outward-open state of a transporter), the simulation can sample transitions between these states. Our group has previously developed and applied such Gō-Martini approaches, including a switching protocol to generate transition pathways and a multiple-basin method to simulate equilibrium fluctuations between states. ^17,18^ The latter employs the exponential mixing scheme (EXP) to create a combined energy landscape smoothly between the basins of two reference conformations.

Despite their theoretical promise, the practical application of Gō-Martini models has been hampered by several technical and methodological barriers. First, in standard Gō-Martini implementations, the Gō-contacts are often implemented as part of the nonbonded potential. This can lead to an over-stabilization of non-specific protein-protein interactions at the surface, promoting spurious aggregation in multi-copy simulations and distorting protein-environment interfaces. ^16,19^ Second, the definition and implementation of mixed topologies that incorporate interactions from two or more reference states are non-trivial and require significant manual intervention, particularly when extending to new interactions. ^18^ This complexity has impeded the exploration and integration of different mixing schemes, such as Hamiltonian-mixing method (HAM). ^20^ Third, the resulting energy landscapes are highly sensitive to parameters that govern the relative stability of the states and the height of the barrier between them. Optimizing these parameters to match experimental observables is often a tedious trial-and-error process.

To address these challenges and provide a robust, accessible platform for the community, in this study we present CTGoMartini (**C**onformational-**T**ransition **G**ō**-Martini**), a comprehensive Python package built on the OpenMM simulation engine and integrated with Martinize2 for automated topology generation. CTGoMartini automates the entire workflow for conformational transition studies: from processing multiple input structures and identifying native contacts, to constructing hybrid topologies, running MD simulations, and analyzing results. A key innovation is the redefinition of the Gō contact potential as a new interaction type, which prevents spurious aggregation artifacts. The package implements both switching and multiple-basin (using EXP or HAM mixing schemes) Gō-Martini protocols. Furthermore, we integrate Hamiltonian replica exchange molecular dynamics (HREMD) ^21,22^ with the PyMBAR package ^23^ to systematically optimize mixing parameters and efficiently estimate free energy surfaces.

We demonstrate the utility of CTGoMartini through two biologically relevant case studies. First, we simulated the lipid transporter Spinster homolog 2 (SPNS2), capturing its transition between inward-open and outward-open states and the coupled translocation process of its sphingosine-1-phosphate (S1P) substrate. Second, we performed a systematic investigation of the mechanosensitive ion channel TREK1, elucidating how membrane surface tension and specific anionic lipids (POPA and PIP_2_) modulate its conformational equilibrium. These examples showcase the package’s capability to provide mechanistic insights into complex lipid-involved conformational transitions that are difficult to access with other computational methods.

## Results

### Overview of the CTGoMartini Framework

The CTGoMartini package is a Python-based framework for constructing, executing, and analyzing protein conformational transitions using Gō-Martini models. Built on the Martinize2 topology builder ^24^ and the OpenMM simulation engine, ^25^ the framework streamlines the entire workflow, from topology generation, to performing simulations and analyzing transition pathways. As illustrated in Fig. 1a, this workflow begins with processing the atomic structures of two distinct conformational states of a protein to generate contact maps, which are then converted into Gō-Martini topologies. CTGoMartini subsequently generates appropriate mixing topologies for two primary simulation approaches: the switching method, which uses two separate topologies, and the multiple-basin method, which combines both states into a mixed topology. The package interfaces with OpenMM to run the simulations, followed by analysis of the resulting conformational transitions.

**Figure 1:**
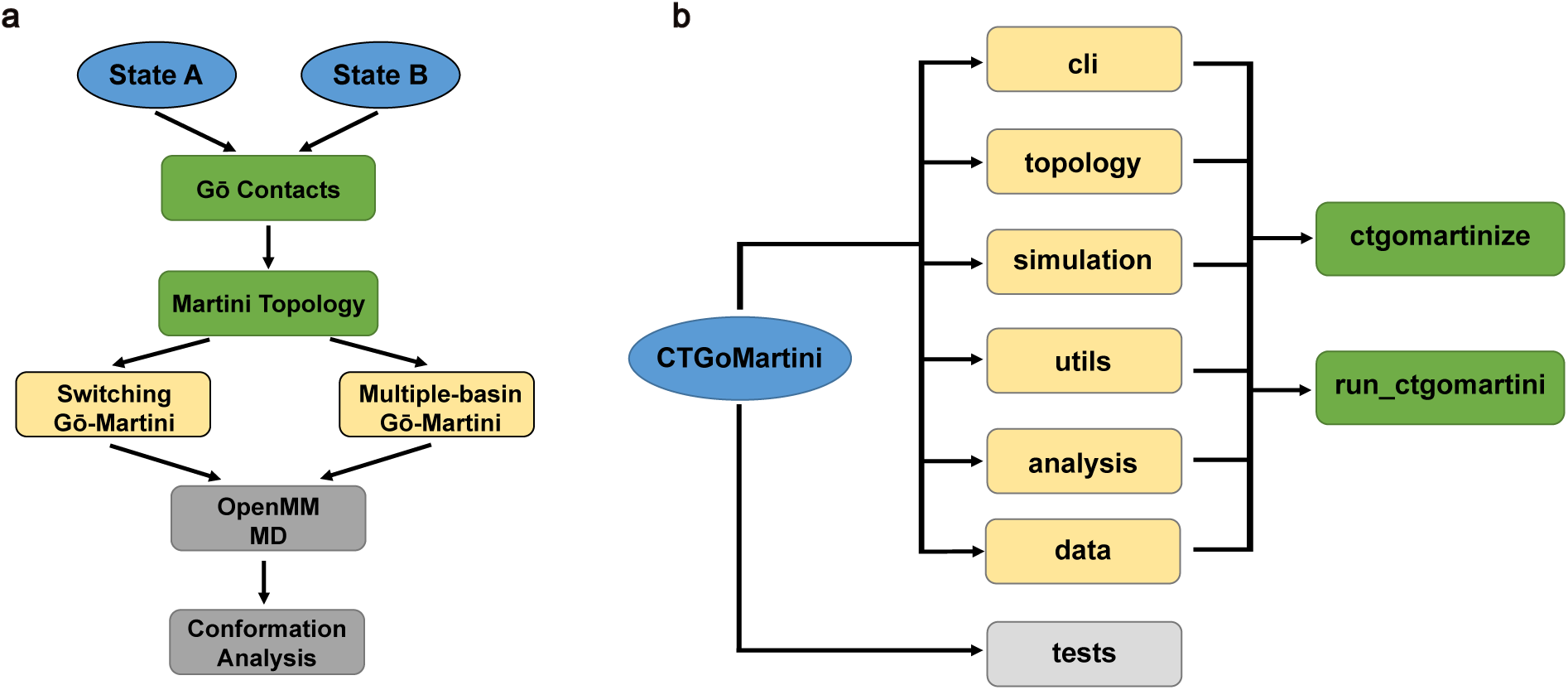
Workflow and architecture of the CTGoMartini framework. **(a)** Schematic workflow of CTGoMartini for protein conformational transitions. (1) Atomic structures of two protein states are processed to generate contact maps; (2) Gō-Martini topologies are built using Martinize2; (3) CTGoMartini prepares separate topologies for the switching method or generates mixed topologies for the multiple-basin method; (4) Simulations are performed using OpenMM; (5) Resulting conformational transitions are analyzed. **(b)** Software architecture diagram illustrating the core modules and their interactions.

CTGoMartini is organized into six core modules (Fig. 1b): topology (parsing GROMACS-style topology files and building OpenMM systems), simulation (managing simulation execution), cli (providing command-line interfaces), utils (handling structure I/O operations and helper functions), analysis (supplying post-processing tools), and data (containing force field parameters and templates). Many comprehensive tests are also included for validation. This modular design ensures extensibility of different interaction types and ease of use, allowing users to set up complex simulations through two main command-line tools: ctgomartinize for topology generation and run_ctgomartini for executing simulations with MD or HREMD modes.

### A Redefined Contact Potential Prevents Spurious Protein Aggregation

In the standard Gō-Martini 3 model, native contact interactions are implemented as standard nonbonded Lennard-Jones potentials projected through virtual sites, with a unified 1.1 nm cut-off. ^16^ Because these interactions are defined solely by bead types, different atoms with identical bead types from different copies of the same protein can interact attractively. This leads to an over-stabilization of non-native protein-protein interactions, resulting in spurious aggregation in multi-copy simulations.

To resolve this, CTGoMartini introduces a redefined Gō contact as a new interaction type named “contacts”. This Gō potential (Gō-Contacts) retains the functional form of the standard Martini nonbonded potential (Gō-Nonbonded) but is explicitly restricted to pairs of beads defined as native contacts within a single protein, thereby eliminating artifactual intermolecular attraction. We first validated the numerical equivalence of this new implementation against the original. Using eight diverse single protein systems, we compared the single-point energies and forces computed by the standard Gō-Nonbonded model in GROMACS with those from the Gō-Contacts model using CTGoMartini in double precision. The results showed excellent agreement that the absolute energy error is below 10^−3^ kJ/mol with relative error < 10^−5^, and the absolute force error is below 10^−4^ kJ/mol/nm with relative error < 10^−5^ (Supplementary Table S1).

We further validated the Gō-Contacts model by simulating multiple copies of three structurally distinct proteins: the transmembrane helix KALP in a lipid bilayer, the soluble Protein G and Ubiquitin. As shown in Fig. 2 and Supplementary Fig. S1, simulations using the standard Gō-Martini (Gō-Nonbonded) model exhibited pronounced, non-physical aggregation, evidenced by a significant reduction in solvent-accessible surface area (SASA) and decreased intermolecular distances. In contrast, both the pure elastic network (Elastic) and the new Gō-Contacts model maintained a well-dispersed state for all proteins. The performance of the Gō-Contacts model was consistent with that of the Gō-Elastic model, confirming that the redefined contact potential successfully eliminates aggregation artifacts while preserving the stabilizing effect of native contacts. Furthermore, this redefined contact potential provides the foundation for constructing a multiple-basin energy landscape, and its separate functional form naturally facilitates such an extension.

**Figure 2:**
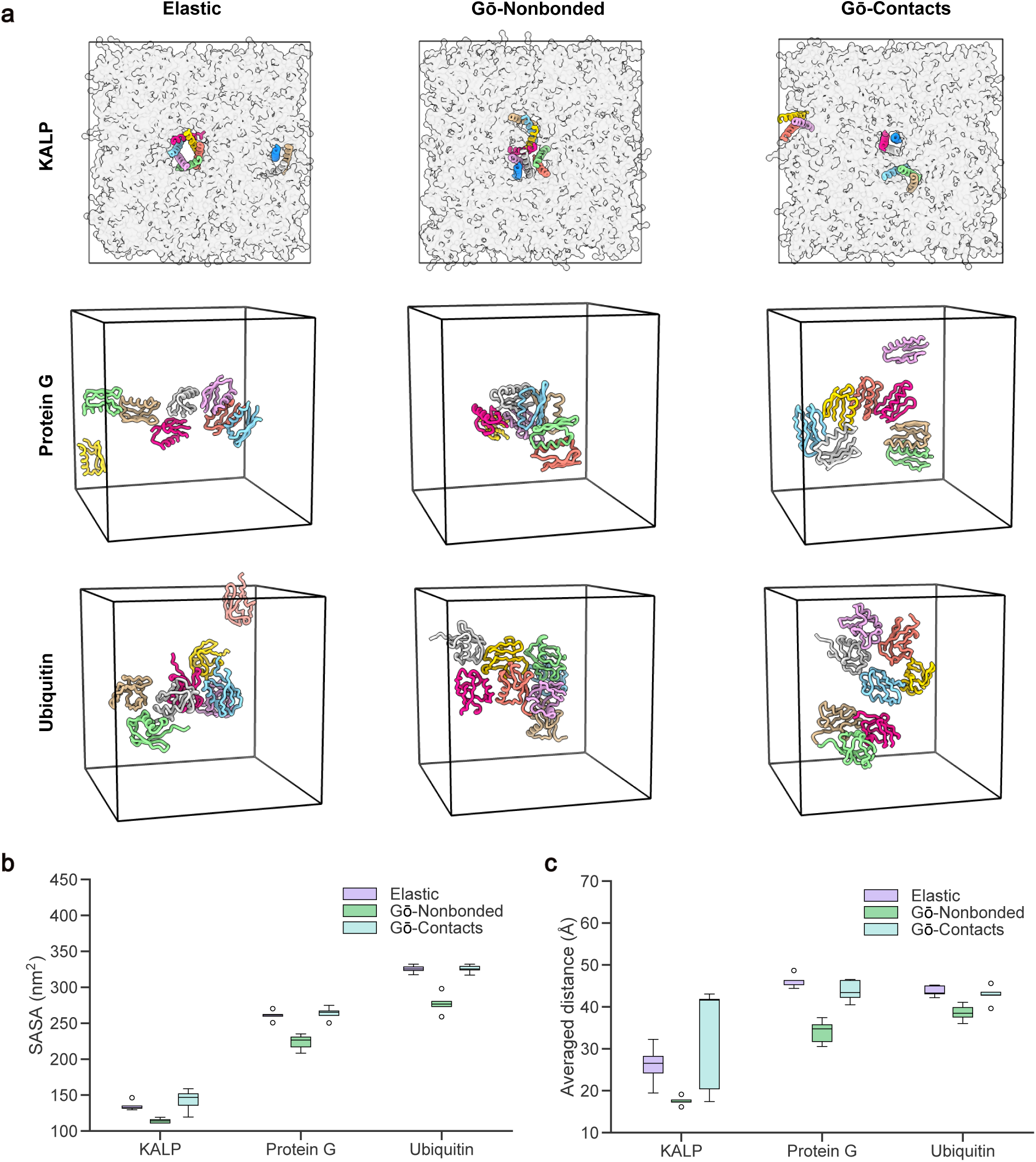
Comparison of protein aggregation behaviors using different Martini models. **a,** Representative snapshots for KALP, Protein G, and Ubiquitin systems using the Elastic, Gō-Nonbonded, and Gō-Contact models, respectively. Proteins are colored individually to distinguish distinct copies. **b,** Box plots showing the SASA for each system across the three models. **c,** Box plots of the averaged intermolecular distance for each system across the three models. Data in **b** and **c** are derived from five independent replicates for each condition, with the last 4000 ns of each trajectory used during the analysis.

### Switching and Multiple-Basin Gō-Martini Methods Enable Conformational Transitions

The CTGoMartini framework implements three complementary methods for sampling conformational transitions in proteins, as illustrated in Fig. 3a. The switching method iteratively alternates the underlying Gō-Martini potentials between two reference states over time, generating a directed transition pathway. In contrast, the multiple-basin methods construct a unified energy landscape that enables spontaneous reversible transitions by mixing two single-basin potentials. CTGoMartini provides two established mixing schemes: EXP and HAM mixing schemes. The EXP scheme approximates a Boltzmann-weighted combination of the two underlying potentials. ^26^ Its energy landscape is shaped by three parameters in which *β* modulates the height of the transition barrier, while *C*_1_ and *C*_2_ tune the relative depths of the two energy basins. The HAM scheme, inspired by quantum mechanical coupling, uses parameters *Δ*, *C*_1_, and *C*_2_ to similarly control the barrier and basin stability. ^20^ Detailed mathematical formulations for both mixing schemes are provided in the Methods section.

**Figure 3:**
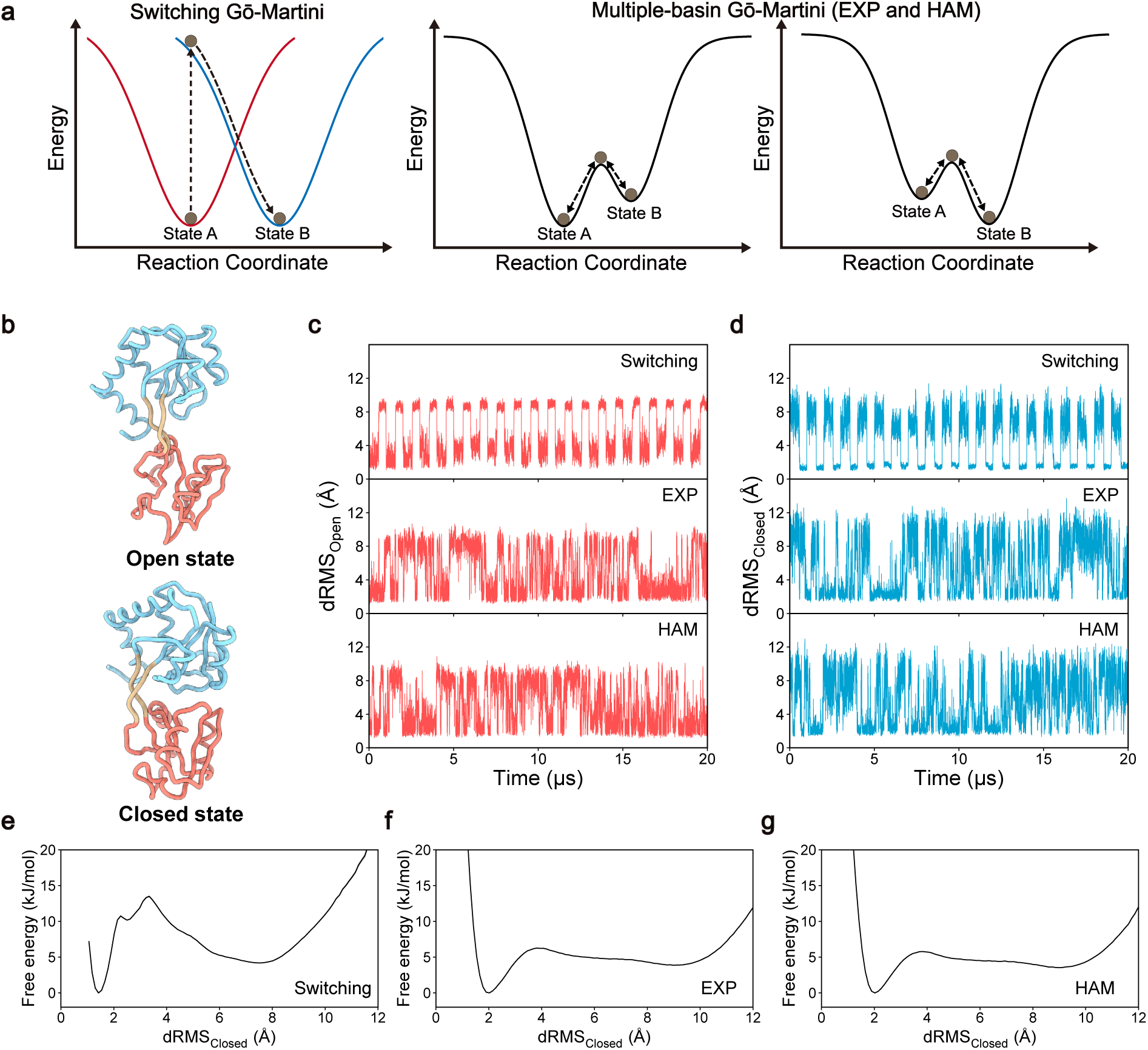
Comparison of Switching and Multiple-basin Gō-Martini approaches for simulating protein conformational transitions. **a,** Schematic illustration of the Switching Gō-Martini and Multiple-basin Gō-Martini methods. **b,** Coarse-grained structures of GlnBP in the open (PDB ID: 1GGG ^28^) and closed (PDB ID: 1WDN ^29^) states. The protein is shown as a ribbon, with the large domain in sky blue, the small domain in salmon, and the hinge region in tan. **c, d,** Time evolution of dRMS with respect to the open state (**c**) and closed state (**d**) over 20 µs simulations. **e-g,** Free energy profiles of GlnBP as a function of dRMS_Closed_ for the Switching (**e**), EXP(**f**), and HAM (**g**) methods, respectively. Profiles are derived from 10 independent replicates, with the first 1 µs of each trajectory discarded during the analysis. Shaded areas represent the standard deviation estimated using bootstrap analysis. Note that these errors are negligibly small and thus not visible in the figure.

The single-basin Gō-Martini potential (Gō-Contacts) used in the switching method has been numerically validated in the previous subsection. To verify the implementation of the multiple-basin potentials, we performed single-point energy and force calculations on ten intermediate conformations for two distinct systems: the soluble protein glutamine-binding protein (GlnBP) and the membrane protein β_2_-adrenergic receptor (β_2_AR). Tests were conducted across a range of mixing parameters for both the EXP and HAM mixing schemes. As shown in Supplementary Fig. S2 and Fig. S3, CTGoMartini maintains excellent numerical agreement with reference calculations for both energy and forces, with relative errors consistently below the predefined tolerance threshold as stated above, confirming the precision of the mixed-potential implementation.

To benchmark these methods, we simulated the well-characterized conformational transition of GlnBP between its open and closed states (Fig. 3b). We quantified the overall conformational change using a distance root-mean-square deviation (dRMS) as a reaction coordinate, which calculated the changes of residue-residue distance referred to one state along the simulations with the difference between two states over 5 Å. ^27^ As shown in Fig. 3c–d, the dRMS time series with respect to the open and closed states clearly demonstrate reversible transitions between open and closed states over 20 µs of simulation with all three approaches. With the Switching method, GlnBP iteratively alternates between states on a ∼1 µs cycle, while the EXP and HAM Multiple-basin methods enable spontaneous transitions. For a meaningful comparison, the parameters were selected to yield similar equilibrium constants, Keq_Closed/Open_, of 0.82 (EXP), 0.72 (HAM), and 0.78 (Switching), respectively. Free energy profiles projected along the dRMS reaction coordinate (Fig. 3e–g) reveal that the EXP and HAM schemes produce a broad and shallow barrier (∼6 kJ/mol) that facilitates reversible crossing, in contrast to the higher and less smooth barrier (13.5 kJ/mol) characteristic of the Switching protocol. The consistency of the free-energy landscapes between the EXP and HAM schemes underscores the similarity of these two mixing approaches. Collectively, these results demonstrate that the CTGoMartini framework can robustly model large-scale conformational transitions between defined states. Successful application to the ribose-binding protein (RBP) further validates the generality of our approaches (Supplementary Fig. S4).

### Hamiltonian Replica Exchange and PyMBAR Enable Efficient Parameter Optimization

Manually tuning the parameters that control the relative stability (*C*_1_ and *C*_2_) and barrier height (*β* for EXP scheme or *Δ* for HAM scheme) of multiple-basin potentials to reproduce experimental observables is a challenging and often tedious process. While the Multistate Bennett Acceptance Ratio (MBAR) method can estimate free energies at unsampled parameter values by reweighting trajectories collected at different ones, its accuracy relies heavily on sufficient overlap between the sampled ensembles. ^27^ To overcome this limitation and facilitate the sampling, CTGoMartini integrates HREMD with PyMBAR into a highly efficient workflow. In this scheme, multiple replicas of the system are simulated concurrently, each using a distinct set of mixing parameters(e.g. *C*_1_, *C*_2_, *β* or *Δ*). At regular intervals, pairs of replicas attempt to exchange their parameter sets according to a Metropolis criterion, thereby promoting efficient diffusion through both conformational space and parameter space and ensuring the necessary overlap for subsequent reweighting (Fig. 4a).

**Figure 4:**
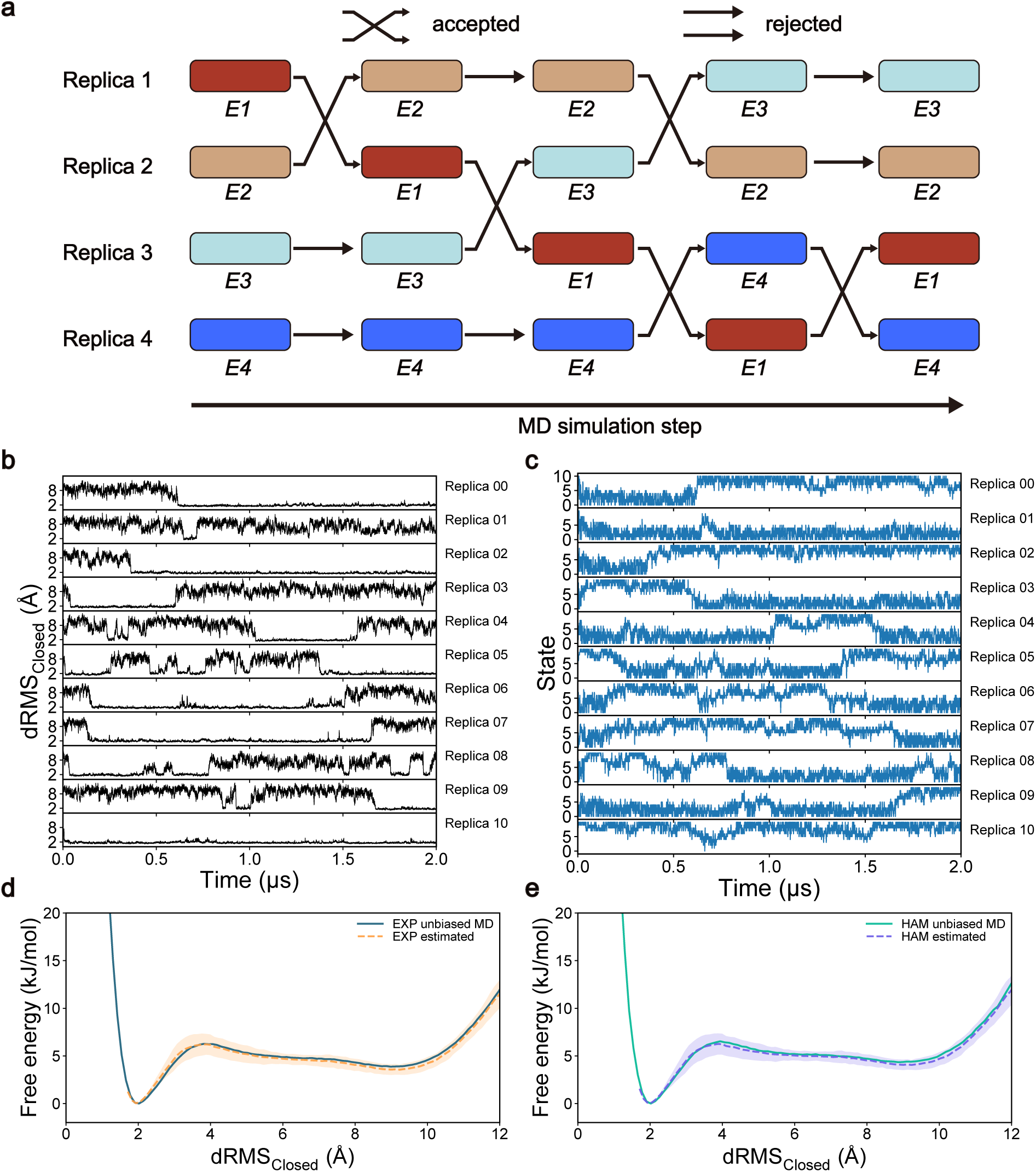
Parameter optimization using Hamiltonian Replica Exchange Molecular Dynamics (HREMD) and PyMBAR. **a,** Schematic illustration of the replica exchange algorithm. Replicas running with different Hamiltonians (represented by colors and labels E1-E4) attempt to swap parameters at periodic intervals. Successful swaps and rejected attempts allow replicas to traverse the parameter space. **b,** Time evolution of the dRMS_Closed_ of GlnBP for 11 individual replicas over a 2 µs simulation. **c,** Time evolution of the Hamiltonian state index of GlnBP for each replica. **d, e,** Comparison of free energy profiles derived from HREMD simulations via PyMBAR predictions (dashed lines) versus long unbiased MD simulations (solid lines). Results are shown for the EXP mixing method (**d**) and the HAM mixing method (**e**). All predictions are derived from PyMBAR using raw HREMD data collected using the EXP mixing method. The shaded regions represent standard deviation calculated from three independent replicates.

We applied this workflow to the open-closed transition of GlnBP using the EXP mixing scheme. An 11-replica HREMD simulation was performed, with *C*_1_ values ranging from −480 to −80 kJ/mol in increments of 40 kJ/mol, intentionally omitting the target value of −300 kJ/mol, while *β* and *C*_2_ were held constant at 1*/*300 mol/kJ and 0 kJ/mol, respectively. The simulation efficiently sampled conformational changes, as evidenced by the fluctuations in dRMS_Closed_ within each replica (Fig. 4b). Concurrently, the frequent exchanges of the Hamiltonian state index demonstrated rapid mixing across the parameter ladder (Fig. 4c).

The combined trajectory data from all replicas were analyzed with PyMBAR to predict equilibrium populations and free energy profiles for any parameter set within the sampled range.

For the deliberately omitted target (*C*_1_ = −300 kJ/mol), the PyMBAR prediction showed excellent agreement with the profile obtained from a long unbiased simulation at the same parameters (Fig. 4d). Remarkably, the same EXP-scheme HREMD data could also be reweighted to accurately predict the profile for the HAM mixing method at its set parameters (*Δ* = 350 kJ/mol, *C*_1_ = −348 kJ/mol, *C*_2_ = 0 kJ/mol) (Fig. 4e), illustrating the consistency of the underlying conformational ensemble and the robustness of the HREMD-PyMBAR approach. We also successfully applied this approach to optimize parameters for RBP, demonstrating its general applicability (Supplementary Fig. S5).

### Investigating Substrate Binding and Translocation in the Lipid Transporter SPNS2

To demonstrate the application of CTGoMartini to complex biological systems, we investigated the transport mechanism of the S1P transporter SPNS2. As a key regulator of immune response, lymphatic function, and vascular integrity, SPNS2 is an important therapeutic target implicated in autoimmune diseases and cancer metastasis. ^30^ While cryo-EM structures of SPNS2 in inward-open and outward-open states have been available, ^31^ fundamental questions regarding the transport cycle remain unresolved. Specifically, the detailed molecular mechanism of S1P translocation across the membrane is unknown and its precise binding pose in the inward-open state is still debated. ^32^

To address these questions, we employed the Switching Gō-Martini method to simulate the conformational transition of SPNS2 from the inward-open to the outward-open state. First, we developed and validated a coarse-grained Martini model for S1P that accurately reproduces its structural and dynamic properties (Supplementary Fig. S6). Using CTGoMartini, we then constructed CG models of the inward-open and outward-open states of SPNS2 (Fig. 5a). Equilibrium simulations were initiated from the inward-open state inserted into a POPC bilayer with S1P placed at its cryo-EM resolved position. From 50 independent replicates (0.5 µs each), S1P remained stably bound in only 13 trajectories, while in the others it dissociated and diffused into the cytoplasmic leaflet (Fig. 5b, Supplementary Fig. S7). In the stable complexes, S1P consistently adopted a distinct binding pose (Pose 1), shifted approximately 9.7 Å deeper into the protein compared to the original coordinates(Fig. 5f and Supplementary Fig. S8). In this pose, the aliphatic tail of S1P inserts into a hydrophobic pocket formed by TM7, TM9, TM10, and TM12, while the phosphate headgroup remains solvent-exposed and interacts with Y120, S242, Y246, and T370. Notably, the Y246A mutation has been reported to significantly reduce SPNS2 transport activity, underscoring the functional relevance of this interaction. ^31^

**Figure 5:**
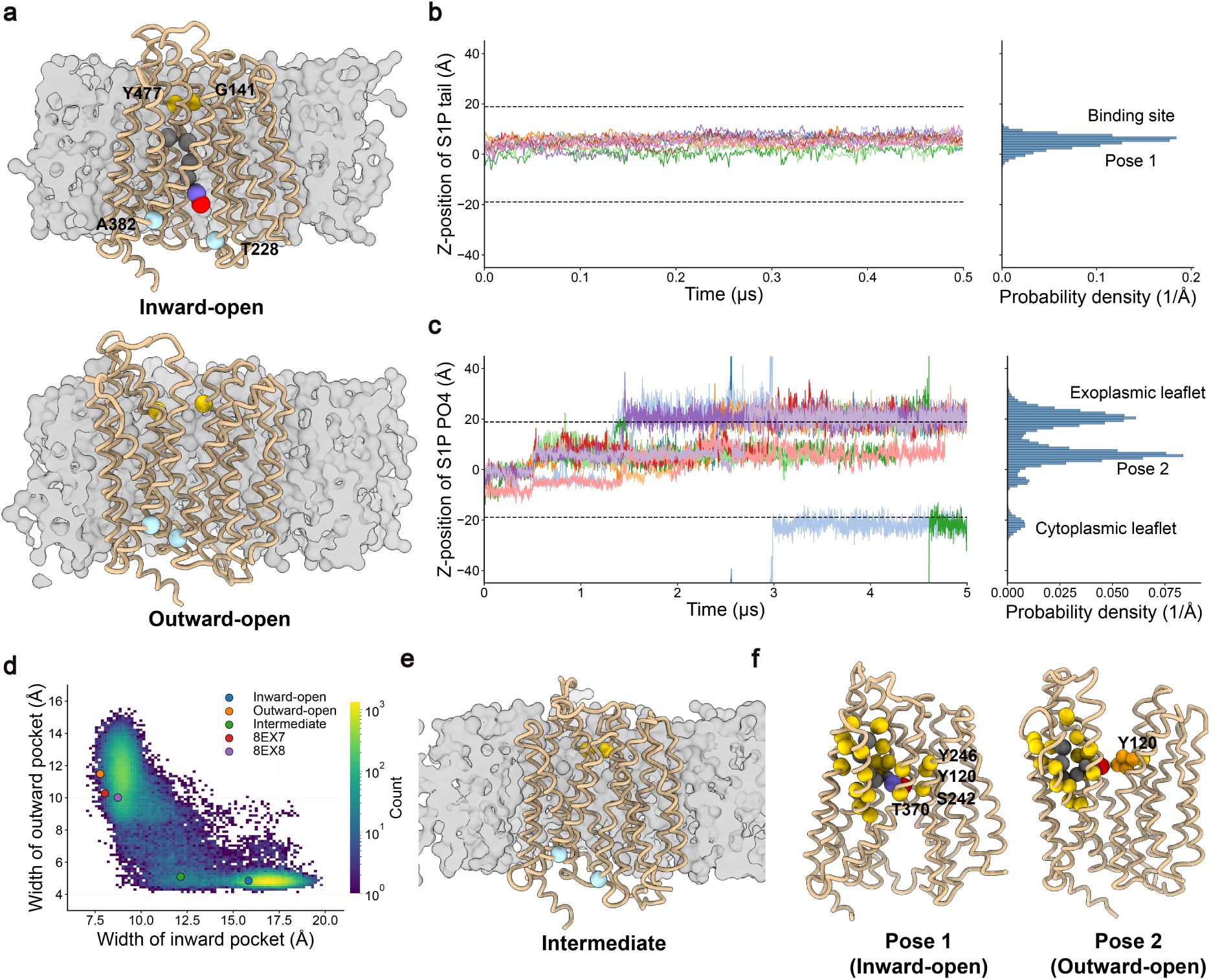
Conformational transitions and substrate translocation process of the SPNS2 transporter. **a,** Coarse-grained structures of SPNS2 in the inward-open and outward-open states (PDB ID: 8EX4 and 8EX5^31^) embedded in a lipid bilayer. The protein is shown as a tan ribbon. Backbone atoms of residues defining width of the outward pocket are highlighted with gold spheres, while those defining the width of the inward pocket are shown as light blue spheres. The membrane is represented by a gray surface. The S1P ligand is depicted within the Inward-open state by using the ball-and-stick model. **b,** Time evolution (left) and probability density distribution (right) of the Z-position of the S1P tail for successful substrate binding processes. **c,** Time evolution (left) and probability density distribution (right) of the Z-position of the S1P phosphate (PO4) group for successful substrate translocation processes. **d,** Two-dimensional histogram of the outward pocket width versus the inward pocket width. The color scale indicates the count of conformations on a logarithmic scale. The circles highlight the coordinates of the representative inward-open, outward-open, intermediate, and the outward-facing partially occluded (8EX7, 8EX8) structures. **e,** Representative structure of the intermediate state identified from the conformational landscape. **f,** Detailed view of the binding site residues (gold spheres) and the S1P ligand in the inward-open and outward-open states. The side chain of Y120 in the right panel was shown as orange spheres.

Starting from these 13 stable complexes, we performed extended switching simulations (4 replicates per structure, 4.5 µs each) to drive the transition toward the outward-open state (Supplementary Fig. S7c). In 10 of the 52 attempts, S1P was successfully translocated to the exoplasmic leaflet alongside the protein conformational change (Fig. 5c). In the remaining simulations, S1P was retained in the binding site (Supplementary Fig. S7b). Interestingly, in two translocated trajectories, S1P exited from the upper leaflet of the membrane into the aqueous phase, diffused across the simulation box, and re-entered the lower membrane leaflet (Fig. 5c). We analyzed the transition pathway using the widths of the inward and outward pockets (defined by BB atom distances T228–A382 and G141–Y477, respectively) as reaction coordinates. The progression clearly follows an alternating-access mechanism, sampling a sequence of inward-open → occluded → outward-open states (Fig. 5d). A two-dimensional histogram of the pocket widths captures this progression, with the reported outward-facing partially occluded structures (PDB IDs: 8EX7, 8EX8^31^) located near the outward-open region. Notably, a tightly occluded intermediate state was identified (inward-pocket width = 12.1 Å, outward-pocket width = 5.1 Å), providing new mechanistic insights into the transport cycle (Fig. 5e).

Simultaneous tracking of the substrate revealed that S1P first changes its binding pose (from Pose 1 to Pose 2) as SPNS2 transitions from the inward-open to the outward-open state, suggesting a stepwise mechanism for S1P translocation (Fig. 5c). In the 10 successful trajectories, S1P subsequently dissociated from the binding pocket to complete transport. The Z-position of the S1P tail shifted only modestly (from 6.5 Å to 9.5 Å in Z-coordinate) between Pose 1 and Pose 2, indicating it remains engaged within the hydrophobic pocket. In contrast, the phosphate headgroup underwent a large displacement (from –1.5 Å to +5.5 Å in Z-coordinate), moving from the intracellular aqueous cavity to the exoplasmic pocket. In the outward-open state, the headgroup of S1P mainly formed an interaction with Y120 (Fig. 5f). The complete translocation process is illustrated in Supplementary Movie S1. These results demonstrate that our simulations provide a detailed mechanistic model for S1P binding and transport by SPNS2, highlighting the capability of CTGoMartini to elucidate complex biomolecular processes that integrate conformational change with ligand dynamics.

### Membrane Tension and Lipid Composition Modulate the Conformational Equilibrium of TREK1

Mechanosensitive two-pore domain potassium (K2P) channels translate membrane-mediated physical and chemical stimuli into electrical signals. TREK1, a prominent member of this family, is regulated by diverse stimuli including membrane stretch, temperature, phosphorylation, and lipids such as phosphatidic acid (PA) and phosphatidylinositol 4,5-bisphosphate (PIP_2_). ^33,34^ A central conformational change involves the transmembrane helix 4 (TM4), which transitions between an “up” state and a “down” state, modulating the lateral fenestrations and ion conductance of channels. Thus, TREK1, as a dimer, can populate three distinct conformational states containing Up/Up, Up/Down, and Down/Down. Recently, all three states of TREK1 structures have been resolved, especially, including a Up/Down apo state and a PA-bound Up/Up conformation. ^35^ In previous work, we used the Multiple-basin Gō-Martini method to capture surface tension-driven conformational changes in the related K2P channel TRAAK. ^18^ However, truncated C-terminal domains in available TRAAK structures limited a detailed analysis of lipid modulation. To overcome this limitation and explore the intricate coupling between membrane properties and channel conformations, we used the CTGoMartini package to study TREK1 which has resolved longer C-terminal helices. We systematically investigated how its conformational equilibrium is modulated by (i) membrane surface tension and (ii) the presence of modulating anionic lipids, thereby providing a more comprehensive view of lipid-dependent conformational transitions in K2P channels.

We first constructed a double-basin energy landscape for TREK1 using the Up/Up and Up/Down states as references (Fig. 6a). The channel was embedded in a pure POPC bilayer, and lateral surface tensions with 0, 5, 10, 15 and 20 mN/m were applied, respectively. Using the dRMS relative to the Up/Up state (dRMS_Up/Up_) as a reaction coordinate, our simulations captured spontaneous transitions between both two states. Representative trajectories (Fig. 6b) show that higher surface tensions progressively shift the conformational ensemble towards the Up/Up state. This shift is quantitatively reflected in the free energy profiles (Fig. 6c). As tension increases from 0 to 20 mN/m, the free energy basin corresponding to the Up/Down state (near dRMS_Up/Up_ = ∼7 Å) rises significantly from −1.2 to 10.9 kJ/mol, indicating the Up/Down state becomes less favorable, while the basin of the Up/Up state is set as 0 for all tensions. By calculating the population fraction of the Up/Up state, we observed a sigmoidal dependence on surface tension (Fig. 6d). Fitting this data with a Boltzmann function yielded a T_50_ (the tension for half Up/Up population) of 8.1 mN/m. This value is in reasonable agreement with experimental estimates of half-maximal activation of 6.4 mN/m, ^36^ especially considering the simplifications of the coarse-grained model and the fact that the metric (Up/Up fraction) is not a direct proxy for ionic current. Representative intermediate structures sampled across all simulated tension ranges are shown in Fig. 6e. These results robustly demonstrate that membrane stretch biases TREK1 towards the Up/Up state.

**Figure 6:**
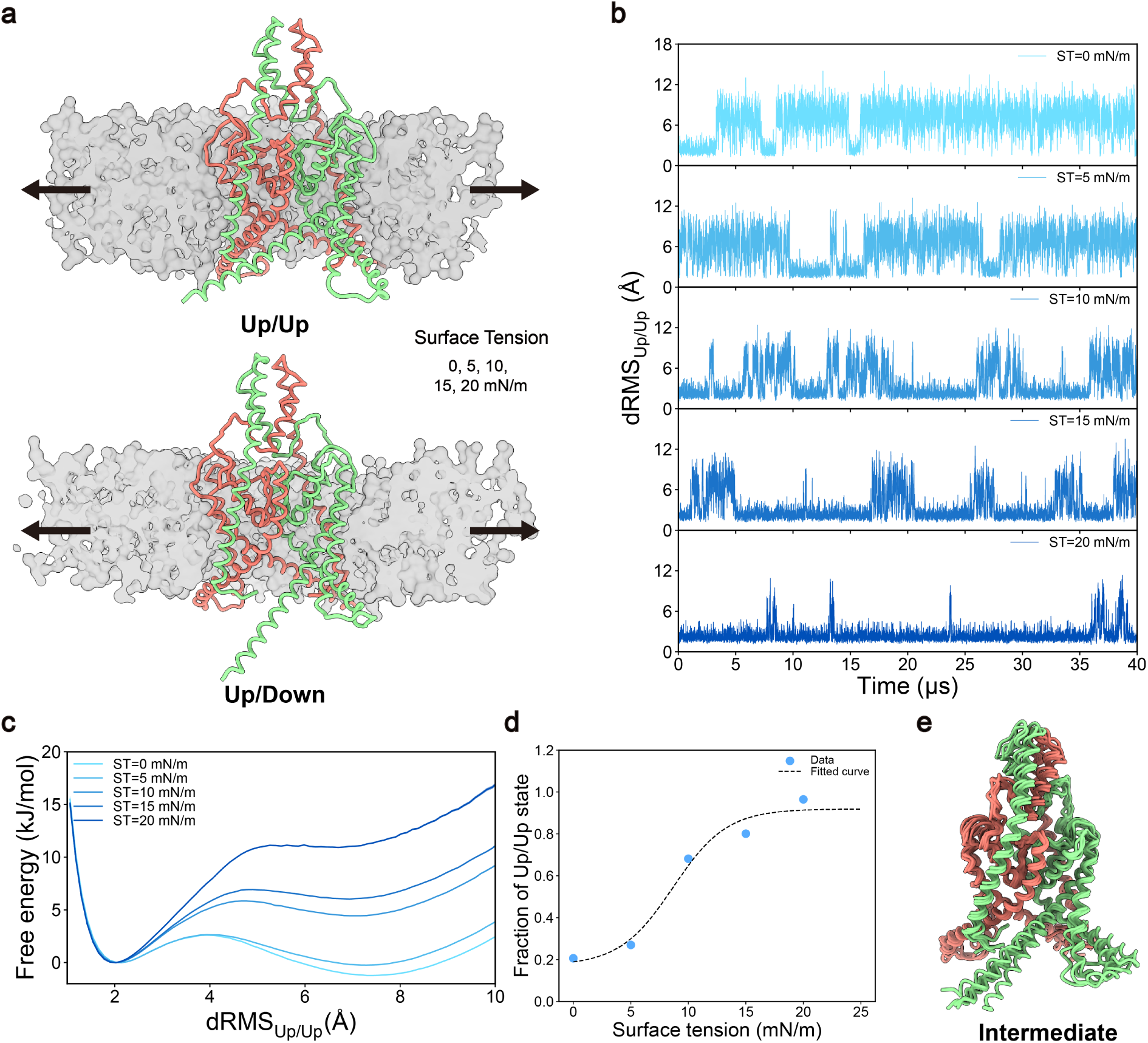
Effect of membrane surface tension on the conformational distribution of TREK1. **a,** Coarse-grained structures of TREK1 in the Up/Up and Up/Down (PDB ID: 8DE8, 8DE7^35^) states embedded in a lipid bilayer. The chains of the protein are colored salmon and light green, respectively. Black arrows indicate the applied lateral tension. **b,** Time evolution of the dRMS with respect to the Up/Up state over simulations. Results are shown for surface tensions ranging from 0 to 20 mN/m. **c,** Free energy profiles projected along the dRMS_Up/Up_ reaction coordinate. Data from all trajectories, excluding the initial 2 µs of each, are used for analysis. The shaded area surrounding each line represents the standard deviation of the free energy, estimated using bootstrap analysis. **d,** The fraction of the Up/Up state population as a function of surface tension. Blue dots represent simulation data, and the dashed line represents a Boltzmann sigmoidal fit. **e,** Representative structures of the intermediate conformational state observed during the transitions for distinct surface tensions.

We next examined the influence of anionic lipids, known biochemical modulators of TREK1. We simulated TREK1 in bilayers containing either 20% POPA or 10% PIP_2_ in the cytoplasmic leaflet with POPC making up the remainder (Fig. 6a), applying a moderate tension of 5 mN/m to better demonstrate lipid effects on the conformational dynamics of TREK1. Compared to pure POPC, both POPA and PIP_2_ markedly stabilized the Up/Up state, as evidenced by the dRMS_Up/Up_ trajectories spending more time near values characteristic of the Up/Up conformation (Fig. 7b). The corresponding free energy profiles (Fig. 7c) show a pronounced stabilization of the Up/Up state and elevated basins of Up/Down state in the presence of these lipids. The population fraction of the Up/Up state increased from approximately 0.27 in pure POPC to 0.44 with POPA and 0.47 with PIP_2_, confirming their role of enhancing the population of the Up/Up state of TREK1. To understand the structural basis of this modulation, we analyzed protein-lipid interaction sites, defining a binding event by a lipid headgroup occupancy exceeding 40%. Both POPA and PIP_2_ headgroups formed stable, state-dependent interaction patterns at the protein-lipid interface (Fig. 7d,e and Supplementary Fig. S10a,c). Residues with a lipid occupancy ratio (Up/Up vs. Up/Down) greater than 1.5 were identified as key anchors that preferentially engage anionic lipids in the Up/Up state (Supplementary Fig. S10b,d). For POPA, these included K304 and R311 on TM4, and T46 on TM1. A similar pattern was observed for PIP_2_, with key interactions at K301, R311, and T46. This suggests a mechanism whereby specific, state-dependent lipid binding acts as a molecular wedge, facilitating and stabilizing the upward movement of TM4. Notably, this polybasic region on TM4 (including R297, K301, K302, K304, and R311) has been experimentally shown to govern TREK1’s sensitivity to PIP_2_, validating our computational findings. ^37^ Collectively, by applying the CTGoMartini method to TREK1, we have quantified two major regulatory inputs that could cooperatively modulate its conformational equilibrium: membrane tension, which directly promotes the Up/Up state, and anionic lipids (POPA, PIP_2_), which stabilize this Up/Up conformation via specific interactions. This study not only reproduces established physiological regulation but also provides an atomistic view of the cooperative interplay between physical tension and chemical lipid composition in governing K2P channel conformational states.

**Figure 7:**
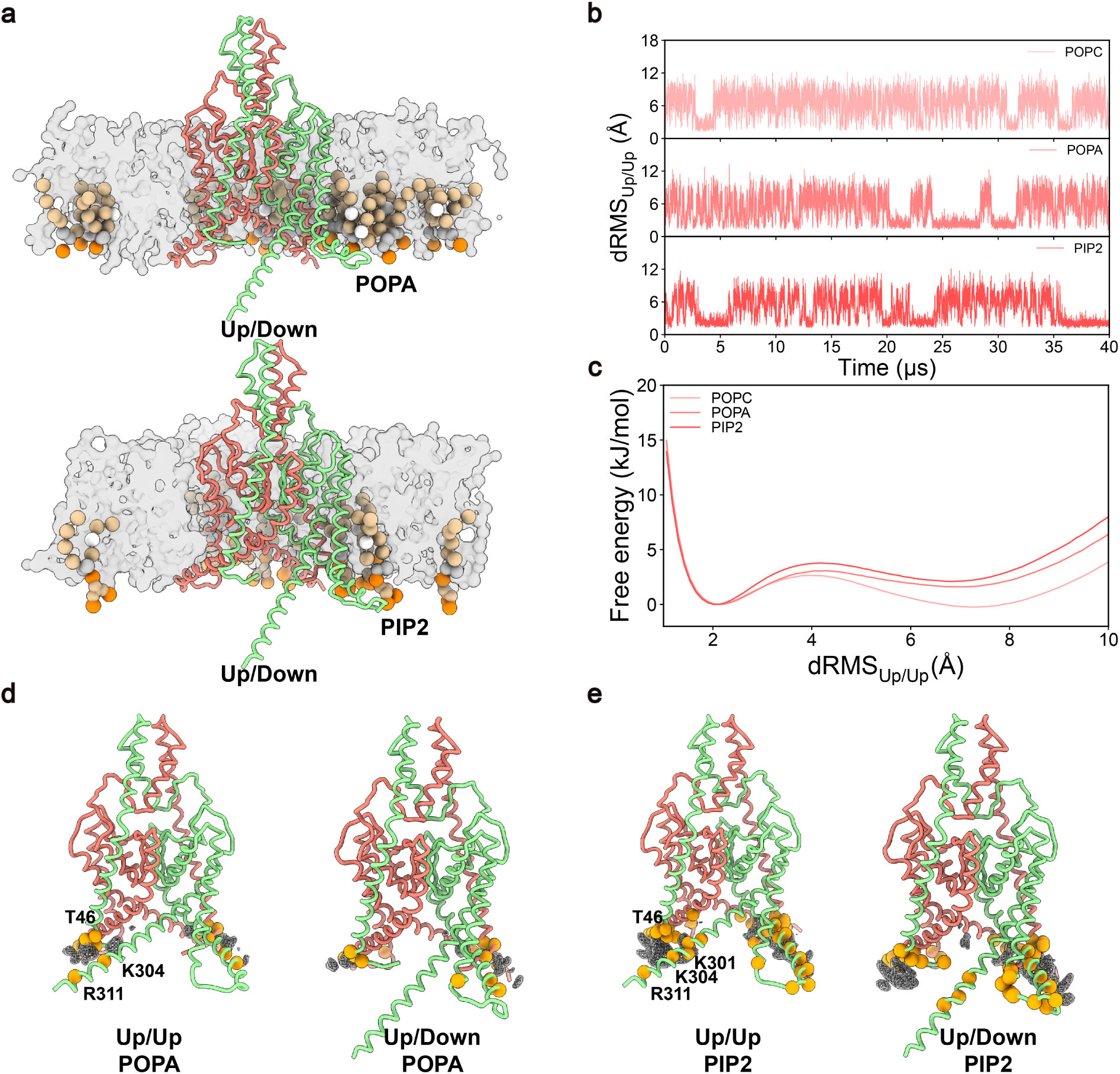
Influence of anionic lipids on the conformational equilibrium of TREK1. **a,** Coarse-grained structures of TREK1 in the Up/Down state embedded in membranes containing POPA (top) and PIP_2_ (bottom). The protein is colored salmon and light green. The specific lipid molecules interacting with the protein are shown as the ball-and-stick model, while the rest of the membrane is gray. **b,** Time evolution of the dRMS with respect to the Up/Up state over simulations. Traces are shown for systems containing pure POPC (top), POPA (middle), and PIP_2_ (bottom). **c,** Free energy profiles projected along the dRMS_Up/Up_ reaction coordinate. Data from all trajectories, excluding the initial 2 µs of each, are used for analysis. The shaded area surrounding each line represents the standard deviation of the free energy, estimated using bootstrap analysis. **d, e,** Detailed structural views of lipid-protein interactions. Key residues with a lipid occupancy greater than 40% are highlighted as orange spheres, which form interactions with POPA (**d**) and PIP_2_ (**e**) headgroups in the Up/Up and Up/Down states, respectively. Lipid binding sites are visualized using a gray mesh representation.

## Discussion

Conformational transitions between distinct structural states are fundamental to protein function, enabling processes such as substrate transport, signal transduction, and ion channel gating. However, elucidating the detailed pathways and environmental modulation of these transitions, particularly in membrane environments, remains a huge challenge. While we have previously developed Switching Gō-Martini and Multiple-basin Gō-Martini methods to address this gap, ^17,18^ their broader application has been hindered by technical barriers and implementation complexity. To facilitate community adoption, here we present CTGoMartini, a comprehensive Python package that implements advanced Gō-Martini methods for simulating protein conformational transitions within explicit membrane environments. CTGoMartini addresses several critical limitations of existing approaches. First, by redefining Gō contacts as a dedicated interaction type rather than embedding them within the nonbonded potential, we eliminate the spurious protein aggregation that hinders traditional Gō-Martini implementations in multi-copy simulations. This innovation ensures accurate representation of protein–protein interfaces, essential for studying multi-protein dynamics. Second, the package provides a unified framework for both nonequilibrium switching simulations and equilibrium multiple-basin simulations, with implementations of both EXP and HAM mixing schemes. Finally, the integration of HREMD with PyMBAR-based analysis automates the traditionally tedious process of parameter optimization, enabling efficient exploration of the energy landscape.

The effectiveness of CTGoMartini is demonstrated through two biologically relevant case studies. For the S1P transporter SPNS2, we captured transitions from inward-open to outward-open states coupled to substrate translocation, identifying a novel intermediate state with a tightly occluded binding pocket. This work provides, to our knowledge, the first dynamic visualization of the S1P translocation process, revealing an intricate single-tail lipid translocation mechanism. For the mechanosensitive ion channel TREK1, we systematically quantified how membrane surface tension and anionic lipid composition (POPA, PIP_2_) modulate the conformational equilibrium between Up/Up and Up/Down states. Our simulations revealed specific lipid-protein interactions that preferentially stabilize particular conformations, offering a rationale for the known modulation of TREK1 by POPA and PIP_2_. These applications highlight CTGoMartini’s unique capability to simultaneously model large-scale protein conformational changes and their coupling to environmental factors, which is a significant advantage over implicit-solvent Gō-models or single-basin Martini simulations.

An unexpected yet functionally relevant insight emerged from the SPNS2 simulations. The substrate binding process critically depended on the protonation state of residue E433. When modeled in its deprotonated (negatively charged) state, S1P failed to bind stably in all 50 independent simulations (Supplementary Fig. S9). In contrast, simulations with E433 protonated (neutral) yielded stable S1P binding (13 out of 50 simulations). This computational observation aligns with experimental reports that mutation of E433 significantly reduces transport activity. ^38^ Additionally, the calculated pKa values for this residue are 7.1 via H++ ^39^ and 7.96 via PROPKA, ^40^ indicating that at physiological pH, E433 likely exists in a dynamic equilibrium between protonated and deprotonated states. Given that SPNS2 functions as a proton-coupled transporter, ^38,41^ the protonation state of E433 may serve as a key element in the transport mechanism, potentially coupling S1P translocation to the proton motive force. This residue could thus act as a proton-sensitive switch, regulating substrate access or release during the alternating-access cycle.

Similarly, our simulations of TREK1 provide deeper insight into the debate regarding PIP_2_ modulation on TREK1. While experimental reports have varied, some suggesting inhibition, ^37,42^ others supporting activation response, ^43,44^ our results clearly show that PIP_2_ stabilizes the Up/Up conformation, which is consistent with recent MD simulation study. ^45^ Although it should be noted that our CG model cannot directly predict ion conductance, the structural shift toward the Up/Up state supports a mechanism in which PIP_2_ binding facilitates the upward movement of TM4, thereby modulating the lateral fenestration and potentially enhancing channel activity. This finding, together with our demonstration that membrane tension also shifts the conformational equilibrium, underscores the utility of CTGoMartini for dissecting complex regulation of membrane proteins.

The computational efficiency of CTGoMartini, which governs its practical applicability, is an important consideration. We therefore conducted a detailed performance comparison (Supplementary Fig. S11). The simulation speed of the single-basin Gō-Nonbonded model in GROMACS is 7.62 µs/day, while that of the single-basin Gō-Contacts model in OpenMM is 7.67 µs/day, demonstrating that our redefined contact implementation achieves comparable performance to the original GROMACS implementation. The multiple-basin Gō-Martini model (EXP mixing scheme) in OpenMM runs at 6.77 µs/day, approximately 12% slower due to the additional overhead of mixing two potentials. Notably, we found that the constraint algorithm in standard Martini models significantly hinders simulation speed. When constraints are replaced by harmonic bonds (with a force constant of 50,000 kJ/mol·nm²), the single-basin Gō-Contacts model achieves 19.7 µs/day, approximately a 2.57-fold increase. Similarly, the multiple-basin model with bonds instead of constraints reaches 12.5 µs/day, a 1.85-fold improvement. Importantly, this conversion has negligible impact on the conformational distribution, as illustrated for GlnBP in Supplementary Fig. S12.

Despite these advances, several limitations warrant consideration. First, like all structure-based models, CTGoMartini requires experimentally determined structures for all conformational states of interest. This prerequisite restricts application to proteins with multiple known structures, though emerging deep-learning methods for predicting alternative conformations may help expand the scope. ^46,47^ Second, the CG representation, although efficient, cannot capture atomic-level details, such as the movement of ions and water in ion channels or the accurate calculation of the associated free energies. Third, the 1.1 nm cutoff for contact interactions, while standard in Martini, truncates longer-range interactions that could be important for certain transitions, particularly those involving large-scale domain rearrangements. ^48,49^ Fourth, the standard Lennard-Jones 6-12 functional form used for the contact potential may not be the optimal interaction type. 10-12 potentials are often used in traditional Gō-like models. ^48,49^ Finally, the removal of side-chain restraints (-scFix) in the multiple-basin scheme, though necessary to avoid steric clashes during transitions, may affect the accuracy of binding-pocket geometries in protein–ligand simulations.

Looking forward, the modular architecture of CTGoMartini is designed for extensibility. It readily accommodates the integration of new mixing schemes, additional contact interaction types, and other enhanced-sampling techniques. Furthermore, the framework can be combined with resolution transformation methods, such as CG2AT2, ^50^ to reconstruct atomistic details along transition pathways, bridging the gap between CG efficiency and all-atom accuracy. We also envision applying this package to systems beyond single protein transitions, including the study of concerted conformational changes in multi-protein assemblies, such as protein-involved membrane fusion and fission. ^51^ Moreover, the CTGoMartini framework can be extended to biomolecular systems beyond proteins, such as RNA ^52^ and DNA, ^53^ and even to general chemical molecules that undergo multiple state conformational transitions. ^54^

In summary, CTGoMartini represents a significant step forward in computational tools for studying protein conformational transitions. By addressing key technical challenges in model implementation and mixing parameter optimization, the package enables detailed investigations of complex biological processes that were previously inaccessible. As an open-source, extensible platform built on widely adopted standards (OpenMM, Martini), CTGoMartini lowers the barrier for researchers to explore protein dynamics in more realistic membrane environments. We anticipate that this tool will facilitate deeper mechanistic understanding of protein functions and contribute to rational design of therapeutics targeting specific conformational states.

## Methods

### Implementation of CTGoMartini

CTGoMartini is built upon the Martini 3 force field, ^13^ which provides coarse-grained representations for proteins, lipids, and small molecules. The package utilizes the OpenMM simulation engine ^25,55^ for all molecular dynamics calculations. For studying protein conformational transitions, CTGoMartini implements three distinct Gō-Martini approaches: (1) single-basin Gō-Martini for stabilizing individual conformational states, (2) Switching Gō-Martini for generating nonequilibrium transition pathways, and (3) Multiple-basin Gō-Martini for simulating spontaneous conformational transitions and investigating environmental modulation on these transitions.

#### Redefined Contact Potential

In standard Gō-Martini implementations, native contacts are typically encoded within the nonbonded potential, which can lead to spurious aggregation artifacts due to mistaken proteinprotein interactions between multiple copies of the same protein.To address this limitation, CTGoMartini redefines the Gō contact potential as a distinct interaction category named *contacts*, implemented through OpenMM’s CustomBondForce. This potential employs a cutoff-style 6-12 Lennard-Jones functional form with a cutoff distance of 1.1 nm, maintaining the same interaction energy between contacting residues as in standard Gō-Martini.

#### Multiple-Basin Mixing Schemes

For simulations of conformational transitions between two states (State 1 and State 2), CTGo-Martini implements two mixing schemes: ^20,26^

**Exponential (EXP) Mixing Scheme:**

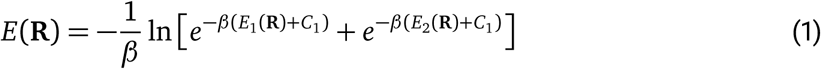

#### Hamiltonian (HAM) Mixing Scheme

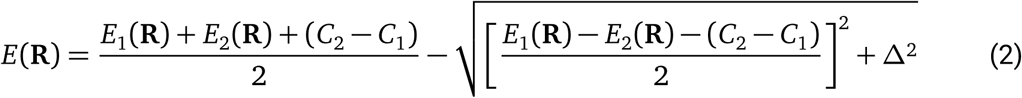

where *E*_1_ and *E*_2_ represent the single-basin potentials for states 1 and 2, respectively; *E* denotes the resulting multiple-basin potential; *C*_1_ and *C*_2_ are constants that control the relative stability of the two states; *β* or *Δ* is a mixing parameter that controls the barrier height between states; **R** is the coordinates of the protein. These mixed potentials are implemented using OpenMM’s CustomCVForce framework, enabling efficient potential mixing according to the specified formulas.

#### Bond, Angle, and Dihedral Treatments in Multiple-basin Mixing Schemes

Standard Martini bonded terms (bonds, angles, and dihedrals) are retained from the force field for single-basin Gō-Martini simulations, with modifications for multiple-basin implementations. To reduce computational cost, bonds, angles, and dihedrals with equilibrium values within a specified cutoff between the two reference states are combined and averaged based on the specific combination rules. ^18^ Side chain restraints (implemented via the *-scfix* flag in Martinize2) are maintained in the standard single-basin Martini model but are discarded in the multiple-basin scheme to allow side chain rearrangements during conformational transitions.

#### Validation via Single-Point Energy and Force Calculations

To validate the numerical accuracy of our implementation, we performed single-point energy and force calculations for both single-basin and multiple-basin Gō-Martini models. For the single-basin Gō-Martini potential, energy and force evaluations were conducted using both the CTGo-Martini (OpenMM) implementation and a reference GROMACS implementation for eight protein systems spanning diverse structural classes (Supplementary Table S1). For energy comparison, the total potential energy was extracted from each platform. Adopting the validation criteria from the Martini-OpenMM implementation, ^55^ we considered the implementations to be in agreement if the absolute energy error was below 10^−3^ kJ/mol and the relative error was below 10^−5^.

For force validation, the force vectors on each atom, **F***_i_*, were extracted from both implementations, excluding forces on virtual sites by setting them to zero prior to analysis. Agreement was assessed using a combined absolute and relative tolerance criterion for each atom *i*. For a pair of force vectors **F**^(OpenMM)^ and **F**^(GROMACS)^, the condition for acceptance is:

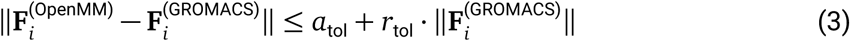

with absolute tolerance *a*_tol_ = 10^−4^ kJ/mol/nm and relative tolerance *r*_tol_ = 10^−5^. The implementation was deemed valid only if this condition held for all atoms. For reporting, the atom with the largest deviation from this inequality was identified, and its absolute and relative errors represent the worst discrepancy in the system. All single-point calculations were performed using GROMACS compiled in double precision and OpenMM in double-precision mode on the Reference platform to minimize numerical differences.

For the multiple-basin Gō-Martini models, reference energies and forces were computed by first calculating the single-basin potentials *E*_1_(**R**) and *E*_2_(**R**) and corresponding forces **f**_1_(**r_i_**) and **f**_2_(**r_i_**) for states 1 and 2 using GROMACS, and then applying the mixing formulas. For the EXP mixing scheme, the reference total force on atom *i* is:

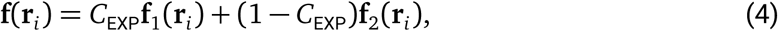

where the mixing coefficient *C*_EXP_ is given by:

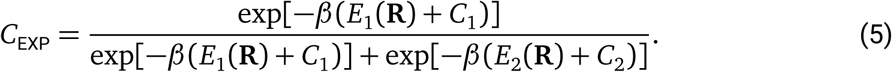

Similarly, for the HAM mixing scheme, the reference total force is:

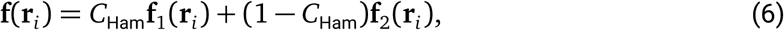

with the mixing coefficient *C*_Ham_ defined as:

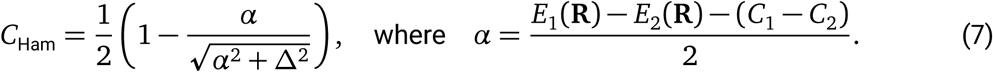

These manually computed reference values were then compared against the energies and forces directly evaluated by CTGoMartini’s multiple-basin implementation. The same absolute and relative tolerance criteria used for the single-basin validation were applied to confirm numerical agreement, ensuring the correct implementation of the mixed potentials in the OpenMM framework.

### General Simulation Settings

All simulations were performed using the Martini 3 coarse-grained force field (version 3.0.0). ^13^ Details regarding the generation of coarse-grained protein models and system setup are provided in the following subsections. For each system, energy minimization was performed, followed by equilibration in the NPT ensemble with position restraints (force constant of 1,000 kJ/mol·nm^2^) applied to the backbone of the CG proteins. Production simulations were conducted at a constant temperature of 310 K and a pressure of 1 bar. For membrane-containing systems, a semiisotropic pressure coupling was used, whereas isotropic pressure coupling was applied to systems without a membrane. Simulations performed with the OpenMM molecular dynamics package ^25^ employed a Langevin integrator ^56^ with a friction coefficient of 0.1 ps^−1^ to maintain temperature, and a Monte Carlo barostat ^57,58^ (semi-isotropic for membranes, isotropic otherwise) with a coupling frequency of 100 steps to maintain pressure. Simulations performed in GROMACS utilized the V-rescale thermostat ^59^ and the Parrinello-Rahman barostat ^60^ to control temperature and pressure, respectively. Both electrostatic and van der Waals interactions were truncated at a cutoff distance of 1.1 nm. Long-range electrostatic interactions were treated using the reaction field method ^61^ with a relative dielectric constant of *ɛ_r_* = 15. During equilibration, a time step of 10 fs was used, while production simulations employed a time step of 20 fs. Unless otherwise specified, simulations were performed using single precision.

### Martini Models and System Setup

#### General Settings for Martini Models

The Martini models used in this study include: (1) the classic single-basin Elastic Network model, (2) the classic single-basin Gō-Martini model, (3) the single-basin Gō-Martini model with redefined contacts, and (4) multiple-basin Gō-Martini models. For the single-basin Elastic model and the single-basin Gō-Martini model, topology parameters were generated using Martinize2. In the Elastic Network model, a force constant of 700 kJ/mol·nm² and a cutoff of 0.5-0.9 nm were applied. The single-basin Gō-Martini model with redefined contacts and the multiple-basin Gō-Martini models were generated using CTGoMartini, which builds upon Martinize2. For all Gō-Martini models, the contact map was calculated based on the Overlap (OV) and repulsive Contacts of Structural Units (rCSU) method. ^62^ These contacts were defined with a distance cutoff between 0.3 and 1.1 nm and a dissociation energy of *ε* = 12.0 kJ/mol. In the multiple-basin Gō-Martini model, to facilitate protein conformational transitions, the long-range external constraints applied to the secondary structure of β-sheets were also converted into contact interactions, while short-range constraints were excluded to avoid redundant interactions in the Gō-like potential.

### Single Point Energy and Force Calculation Systems

To validate the numerical accuracy of the redefined contact potentials and the multiple-basin mixing schemes implemented in CTGoMartini, we performed single-point energy and force calculations on a diverse set of systems. For validating the single-basin Gō-Martini potential, we selected eight proteins spanning various structural classes, including soluble proteins and membrane proteins, which were used in prior Gō-Martini studies ^17,18^ or used in the followed sections. These systems are listed in Supplementary Table S1. For validating the multiple-basin potentials, we used two additional protein systems (GlnBP and β_2_AR) for which well-defined conformational transitions are characterized. For each of these two systems, ten intermediate conformations were selected along previously computed transition pathways ^17^ to test the energy and force evaluation across a range of structures connecting the two reference states.

#### Protein Aggregation Systems

To assess the aggregation behavior of different Martini models, we simulated multiple copies of three representative proteins: KALP (transmembrane α-helix, obtained from MARTINI tutorial), Protein G (soluble protein, PDB ID: 1GB1^63^), and Ubiquitin (soluble protein, PDB ID: 1UBQ ^64^). For each protein, we constructed three distinct models: (1) Standard Martini models with an Elastic Network, (2) Classic Gō-Martini models with native contacts implemented via the nonbonded potential, and (3) Gō-Martini models with redefined contact potential. Models 1 and 2 were generated using Martinize2 and simulated with GROMACS, while model 3 was generated and simulated using CTGoMartini with OpenMM. Each system was solvated in explicit water with 150 mM NaCl using the insane.py script. ^65^ For KALP, peptides were also inserted in the POPC bilayer. 9 copies of KALP, 8 copies of Protein G, and 8 copies of Ubiquitin were evenly distributed in the simulation box. For each condition, we performed five independent 10 µs simulations, and the last 4 µs of each trajectory were used for analysis.

#### GlnBP and RBP

The coarse-grained models of GlnBP were constructed using the Martini 3 force field, based on the open-state (PDB: 1GGG) ^28^ and closed-state (PDB: 1WDN) ^29^ crystal structures. The protein sequences were aligned and trimmed to an identical length (residues 5–224). Topologies for both states were generated as described in the previous section, and multiple-basin Gō-Martini models were constructed by combining the open- and closed-state single-basin models using the specified combination rules. The initial conformation for all simulations was the open state of GlnBP. The protein was solvated in a cubic box (8 × 8 × 8 nm³) using insane.py, ^65^ and 150 mM NaCl was added to maintain charge neutrality. For the switching simulations, the topology of GlnBP was iteratively alternated between the open and closed states, with 556 ns spent in the open state and 444 ns in the closed state per cycle. A 2 ps relaxation interval (1000 steps with a 2 fs time step) was applied between each switch. For the multiple-basin simulations, EXP and HAM mixing schemes were employed. The EXP simulations used parameters *β* = 1*/*300 mol/kJ, *C*_1_ = −300 kJ/mol, and *C*_2_ = 0 kJ/mol, while the HAM simulations used *Δ* = 350 kJ/mol, *C*_1_ = −348 kJ/mol, and *C*_2_ = 0 kJ/mol. Each condition was simulated for 20 µs with 10 independent replicates.

A similar setup was used for the RBP, with open- and closed-state models based on PDB entries 1BA2^66^ and 2DRI, ^67^ respectively. In the switching simulations, RBP was alternated between states with 500 ns per state per cycle, using the same relaxation interval. For multiple-basin simulations, the EXP parameters were *β* = 1*/*175 mol/kJ, *C*_1_ = −198 kJ/mol, *C*_2_ = 0 kJ/mol, and the HAM parameters were *Δ* = 116 kJ/mol, *C*_1_ = −175 kJ/mol, *C*_2_ = 0 kJ/mol. Each condition was simulated for 20 µs with 10 independent replicates.

#### SPNS2

The inward-open and outward-open CG models of SPNS2 were built using the Martini 3 force field, based on structures with PDB codes 8EX4 and 8EX5. ^31^ Missing loops and mutations were repaired using MODELLER ^68^ and the sequences of both proteins were trimmed to have the same length (residues 99-539). Topologies of both states of SPNS2 were generated by CTGoMartini with the switching option. All simulations were initiated from the inward-open conformation. Note that GLU433 of the protein should be protonated to be GLUP, otherwise S1P cannot bind to SPNS2. The protein was then embedded in a POPC bilayer within a rectangular box (8.5 × 8.5 × 10 nm^3^) filled with water using the program insane.py. ^65^ 150 mM NaCl was added to keep the charge of the system neutral. The S1P ligand was parameterized following the Martini 3 small molecule protocol, with atom types assigned based on Martini lipid parameters ^69^ and validated against all-atom simulations (Charmm36m force field). S1P was initially positioned in the binding site of the inward-open state as observed in the cryo-EM structures. The simulation process consisted of an initial 500 ns simulation in the inward-open state across 50 independent replicates. Trajectories in which S1P remained stably bound were selected and used to initiate four parallel simulations each. These systems underwent a brief relaxation (1000 steps with a 2 fs time step), followed by a 4500 ns production simulation in the outward-open state.

#### TREK1

Coarse-grained models of TREK1 in the Up/Up and Up/Down states were constructed using the Martini 3 force field, based on the cryo-EM structures (PDB: 8DE8 and 8DE7^35^). Mutations and missing loops were modeled using MODELLER, ^68^ and the sequences were trimmed to the same length (residues 40-315 for chains A and B). Topologies for both states were generated using CTGoMartini with the EXP mixing option. To accelerate simulation speed, the constraints applied between atoms in the initial structure were converted into harmonic bonds with a force constant of 50,000 kJ/mol·nm². All simulations were initiated from the Up/Up state conformation. The protein was embedded in a symmetric POPC bilayer (147 lipids in the upper and lower leaflets) within a rectangular box (11 × 11 × 13 nm³) and solvated using insane.py. ^65^ The system was neutralized with 150 mM NaCl. Following parameter optimization, the final EXP mixing parameters were set to *β* = 1*/*900 mol/kJ, *C*_1_ = −300 kJ/mol, and *C*_2_ = 0 kJ/mol.

To investigate the influence of membrane surface tension on the conformational distribution of TREK1, a series of simulations were performed at constant lateral tensions of 0, 5, 10, 15, and 20 mN/m. For each tension value, 10 independent simulations of 40 µs each were conducted. To examine the effect of anionic lipids, two additional membrane compositions were prepared: (1) a bilayer with 20% POPA in the lower leaflet, and (2) a bilayer with 10% PIP2 in the lower leaflet. To better present the modulation of lipid composition on TREK1, simulations were performed at a surface tension of 5 mN/m, where small fluctuations induce pronounced shifts in conformational distribution. For each lipid composition, 10 independent 40 µs simulations were performed.

### Hamiltonian Replica Exchange Molecular Dynamics

To enhance sampling efficiency and optimize mixing parameters in multiple-basin simulations, we implemented HREMD within CTGoMartini by using the package OpenMMTools. For each system, 11 replicas were constructed with different parameter sets spanning a range of relative stabilities between two states. Exchange attempts between neighboring replicas were made every 500 steps, with acceptance probabilities calculated using the Metropolis criterion:

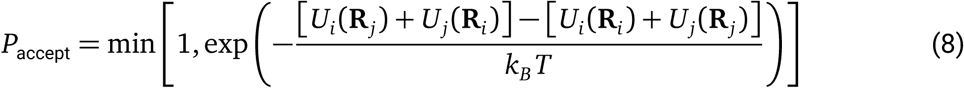

where *U_i_*and *U_j_* denote the potential energies of replicas *i* and *j*, **R***_i_* and **R***_j_* are their respective configurations, *k*_B_ is Boltzmann’s constant, and *T* is the simulation temperature.

For the GlnBP system, the EXP mixing scheme was employed with *β* = 1*/*300 mol/kJ. The stability offset *C*_1_ was varied from −480 to −80 kJ/mol in steps of 40 kJ/mol, while *C*_2_ was fixed at 0 kJ/mol. For the RBP system, we also used the EXP scheme with *β* = 1*/*175 mol/kJ, and *C*_1_ was varied over a range of −250 to −150 kJ/mol with a step size of 10 kJ/mol. For each system, three independent HREMD simulations were performed. Each replica was simulated for 2 µs, with the first 400 ns discarded as equilibration. The resulting trajectories were analyzed using the PyMBAR package ^23^ to compute free energy profiles and predict suitable parameter sets that yield required conformational sampling between states.

### Analysis Methods

Aggregation behavior was quantified by calculating the solvent-accessible surface area using a probe radius of 0.191 nm and 4800 dots per sphere with the GROMACS tool gmx sasa, and by computing the average intermolecular distance between centers of mass of protein backbones using MDAnalysis. ^70^

Conformational similarity was assessed using the distance root mean square (dRMS) deviation metric, defined as:

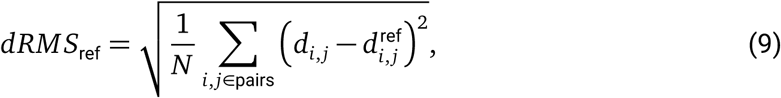

where *d_i_*_,*j*_ and *d*^ref^ represent the distance between the backbone beads of residues *i* and *j* in the sampled conformation and the reference state, respectively. The sum is taken over a set of *N* essential residue pairs, selected according to criteria in which pairs must be separated by at least four residues in sequence and 6–50 Å in space, and must exhibit a distance difference of at least 5 Å between the two reference states.

Free energy profiles along a chosen reaction coordinate *ξ* were calculated from the probability distribution *p*(*ξ*) using:

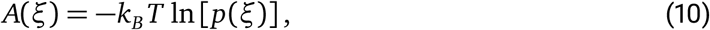

where *A*(*ξ*) is the free energy, *k_B_* is the Boltzmann constant, and *T* is the simulation temperature (310 K). For each simulation, the first 1 µs was discarded as equilibration. For visual comparison, each free energy profile was normalized by subtracting its specific basin, setting the basin value to zero. Standard deviation was estimated using bootstrap analysis with 200 rounds of resampling from the trajectory data.

Lipid–protein interactions were quantified by computing lipid occupancy, defined as the fraction of simulation frames in which the head group or tail group of a lipid molecule was within a cutoff distance of 0.65 nm from any protein bead. To classify simulation frames as belonging to the Up/Up or Up/Down state of TREK1, we used the free-energy profile projected along the dRMS_Up/Up_ reaction coordinate. The position of the barrier separating the two basins was identified, and frames with dRMS_Up/Up_ values lower than the barrier were assigned to the Up/Up state, while frames with higher values were assigned to the Up/Down state.

The representative intermediate structure in SPNS2 was identified by applying the k-means clustering algorithm to the conformational ensemble along the relevant reaction coordinates. All distance calculations and trajectory analysis were performed using MDAnalysis, ^70^ and structural figures were rendered with ChimeraX. ^71^

## Supporting information

SPNS2

## Data Availability

The codes and tutorials for CTGoMartini are available at https://github.com/ComputBiophys/CTGoMartini.

## Acknowledgements

We thank Dr. Xin He at Zhongguancun Academy for helpful discussions regarding the code implementation. This work was supported by the National Key R&D Program of China (2023YFF1204400 and 2024YFA0916800 to C.S.), and the Innovative Research Group Project of the National Natural Science Foundation of China (T2321001 to C.S.). This work was also supported by the research startup fund from Fuzhou University (Grant 0080/511473 to S.Y.). Part of the MD simulations were performed on the Computing Platform of the Center for Life Sciences at Peking University.

## Author Contributions

S.Y. and C.S. conceived the idea and designed the research. S.Y. conducted simulations and analyzed data. S.Y. and C.S. wrote the manuscript. C.S. acquired funding and supervised the work.

## ASSOCIATED CONTENT

The Supporting Information is available. Supplementary Tables and Figures including energy and force validation, protein aggregation tests, parameter optimization, and additional simulation details for RBP, TREK1, and SPNS2 systems (PDF); Translocation process of S1P mediated by the conformational transitions of SPNS2 (Supplementary Movie 1) (mp4).

## References

[1] Deng, D.; Xu, C.; Sun, P.; Wu, J.; Yan, C.; Hu, M.; Yan, N. Crystal structure of the human glucose transporter GLUT1. Nature 2014, 510, 121–125.

[2] Rasmussen, S. G. F.; Choi, H.-J.; Rosenbaum, D. M.; Kobilka, T. S.; Thian, F. S.; Edwards, P. C.; Burghammer, M.; Ratnala, V. R. P.; Sanishvili, R.; Fischetti, R. F.; Schertler, G. F. X.; Weis, W. I.; Kobilka, B. K. Crystal structure of the human beta2 adrenergic G-protein-coupled receptor. Nature 2007, 450, 383–387.

[3] Gadsby, D. C.; Vergani, P.; Csanády, L. The ABC protein turned chloride channel whose failure causes cystic fibrosis. Nature 2006, 440, 477–483.

[4] Chiti, F.; Dobson, C. M. Protein Misfolding, Amyloid Formation, and Human Disease: A Summary of Progress Over the Last Decade. Annual Review of Biochemistry 2017, 86, 27–68.

[5] Xie, T.; Saleh, T.; Rossi, P.; Kalodimos, C. G. Conformational states dynamically populated by a kinase determine its function. Science 2020, 370, eabc2754.

[6] Banari, A.; Samanta, A. K.; Munke, A.; Laugks, T.; Bajt, S.; Grünewald, K.; Marlovits, T. C.; Küpper, J.; Maia, F. R. N. C.; Chapman, H. N.; Oberthür, D.; Seuring, C. Advancing time-resolved structural biology: latest strategies in cryo-EM and X-ray crystallography. Nature Methods 2025, 22, 1420–1435.

[7] Jumper, J.; Evans, R.; Pritzel, A.; Green, T.; Figurnov, M.; Ronneberger, O.; Tunyasuvunakool, K.; Bates, R.; Žídek, A.; Potapenko, A.; Bridgland, A.; Meyer, C.; Kohl, S. A.; Ballard, A. J.; Cowie, A.; Romera-Paredes, B.; Nikolov, S.; Jain, R.; Adler, J.; Back, T.; Petersen, S.; Reiman, D.; Clancy, E.; Zielinski, M.; Steinegger, M.; Pacholska, M.; Berghammer, T.; Bodenstein, S.; Silver, D.; Vinyals, O.; Senior, A. W.; Kavukcuoglu, K.; Kohli, P.; Hassabis, D. Highly accurate protein structure prediction with AlphaFold. Nature 2021, 596, 583–589.

[8] Abramson, J.; Adler, J.; Dunger, J.; Evans, R.; Green, T.; Pritzel, A.; Ronneberger, O.; Willmore, L.; Ballard, A. J.; Bambrick, J.; others Accurate structure prediction of biomolecular interactions with AlphaFold 3. Nature 2024, 1–3.

[9] Lewis, S.; Hempel, T.; Jiménez-Luna, J.; Gastegger, M.; Xie, Y.; Foong, A. Y. K.; Satorras, V. G.; Abdin, O.; Veeling, B. S.; Zaporozhets, I.; Chen, Y.; Yang, S.; Foster, A. E.; Schneuing, A.; Nigam, J.; Barbero, F.; Stimper, V.; Campbell, J., Andrew Yim; Lienen, M.; Shi, Y.; Zheng, S.; Schulz, H.; Munir, U.; Sordillo, R.; Tomioka, R.; Clementi, C.; Noé, F. Scalable emulation of protein equilibrium ensembles with generative deep learning. Science 2025, 389, eadv9817.

[10] Lindorff-Larsen, K.; Piana, S.; Dror, R. O.; Shaw, D. E. How fast-folding proteins fold. Science 2011, 334, 517–520.

[11] Köpfer, D. A.; Song, C.; Gruene, T.; Sheldrick, G. M.; Zachariae, U.; Groot, B. L. D. Ion permeation in K^+^ channels occurs by direct Coulomb knock-on. Science 2014, 346, 352–355.

[12] Kmiecik, S.; Gront, D.; Kolinski, M.; Wieteska, L.; Dawid, A. E.; Kolinski, A. Coarse-grained protein models and their applications. Chemical Reviews 2016, 116, 7898–7936.

[13] Souza, P. C.; Alessandri, R.; Barnoud, J.; Thallmair, S.; Faustino, I.; Grünewald, F.; Patmanidis, I.; Abdizadeh, H.; Bruininks, B. M.; Wassenaar, T. A.; Kroon, P. C.; Melcr, J.; Nieto, V.; Corradi, V.; Khan, H. M.; Domański, J.; Javanainen, M.; Martinez-Seara, H.; Reuter, N.; Best, R. B.; Vattulainen, I.; Monticelli, L.; Periole, X.; Tieleman, D. P.; de Vries, A. H.; Marrink, S. J. Martini 3: A general purpose force field for coarse-grained molecular dynamics. Nature Methods 2021, 18, 382–388.

[14] Periole, X.; Cavalli, M.; Marrink, S.-J.; Ceruso, M. A. Combining an elastic network with a coarse-grained molecular force field: Structure, dynamics, and intermolecular recognition. Journal of Chemical Theory and Computation 2009, 5, 2531–2543.

[15] Poma, A. B.; Cieplak, M.; Theodorakis, P. E. Combining the MARTINI and structure-based coarse-grained approaches for the molecular dynamics studies of conformational transitions in proteins. Journal of Chemical Theory and Computation 2017, 13, 1366–1374.

[16] Souza, P. C. T.; Borges-Araújo, L.; Brasnett, C.; Moreira, R. A.; Grünewald, F.; Park, P.; Wang, L.; Razmazma, H.; Borges-Araújo, A. C.; Cofas-Vargas, L. F.; Monticelli, L.; Mera-Adasme, R.; Melo, M. N.; Wu, S.; Marrink, S. J.; Poma, A. B.; Thallmair, S. GōMartini 3: From large conformational changes in proteins to environmental bias corrections. Nature Communications 2025, 16, 4051.

[17] Yang, S.; Song, C. Switching Gō-Martini for investigating protein conformational transitions and associated protein–lipid interactions. Journal of Chemical Theory and Computation 2024, 20, 2618–2629.

[18] Yang, S.; Song, C. Multiple-Basin Gō-Martini for Investigating Conformational Transitions and Environmental Interactions of Proteins. Journal of Chemical Theory and Computation 2025, 21, 5304–5321.

[19] Korshunova, K.; Kiuru, J.; Liekkinen, J.; Enkavi, G.; Vattulainen, I.; Bruininks, B. M. H. Martini 3 OliGomers: A scalable approach for multimers and fibrils in GROMACS. Journal of Chemical Theory and Computation 2024, 20, 7635–7645.

[20] Okazaki, K. I.; Koga, N.; Takada, S.; Onuchic, J. N.; Wolynes, P. G. Multiple-basin energy landscapes for large-amplitude conformational motions of proteins: Structure-based molecular dynamics simulations. Proceedings of the National Academy of Sciences of the United States of America 2006, 103, 11844–11849.

[21] Sugita, Y.; Kitao, A.; Okamoto, Y. Multidimensional replica-exchange method for free-energy calculations. The Journal of Chemical Physics 2000, 113, 6042–6051.

[22] Fukunishi, H.; Watanabe, O.; Takada, S. On the Hamiltonian replica exchange method for efficient sampling of biomolecular systems: Application to protein structure prediction. The Journal of Chemical Physics 2002, 116, 9058–9067.

[23] Shirts, M. R.; Chodera, J. D. Statistically optimal analysis of samples from multiple equilibrium states. The Journal of Chemical Physics 2008, 129, 124105.

[24] Kroon, P. C.; Grünewald, F.; Barnoud, J.; van Tilburg, M.; Brasnett, C.; Souza, P. C.; Wassenaar, T. A.; Marrink, S. J. Martinize2 and Vermouth provide a unified framework for molecular topology generation. eLife 2025, 12, RP90627.

[25] Eastman, P.; Galvelis, R.; Peláez, R. P.; Abreu, C. R.; Farr, S. E.; Gallicchio, E.; Gorenko, A.; Henry, M. M.; Hu, F.; Huang, J.; others OpenMM 8: molecular dynamics simulation with machine learning potentials. The Journal of Physical Chemistry B 2023, 128, 109–116.

[26] Best, R. B.; Chen, Y. G.; Hummer, G. Slow protein conformational dynamics from multiple experimental structures: The helix/sheet transition of Arc repressor. Structure 2005, 13, 1755–1763.

[27] Shinobu, A.; Kobayashi, C.; Matsunaga, Y.; Sugita, Y. Building a macro-mixing dual-basin Go model using the multistate Bennett acceptance ratio. Biophysical Journal 2020, 118, 179a.

[28] Hsiao, C. D.; Sun, Y. J.; Rose, J.; Wang, B. C. The crystal structure of glutamine-binding protein from Escherichia coli. Journal of Molecular Biology 1996, 262, 225–242.

[29] Sun, Y. J.; Rose, J.; Wang, B. C.; Hsiao, C. D. The structure of glutamine-binding protein complexed with glutamine at 1.94 Å resolution: Comparisons with other amino acid binding proteins. Journal of Molecular Biology 1998, 278, 219–229.

[30] Spiegel, S.; Maczis, M. A.; Maceyka, M.; Milstien, S. New insights into functions of the sphingosine-1-phosphate transporter SPNS2. Journal of Lipid Research 2019, 60, 484–489.

[31] Chen, H.; Ahmed, S.; Zhao, H.; Elghobashi-Meinhardt, N.; Dai, Y.; Kim, J. H.; McDonald, J. G.; Li, X.; Lee, C.-H. Structural and functional insights into Spns2-mediated transport of sphingosine-1-phosphate. Cell 2023, 186, 2644–2655.e16.

[32] Li, H. Z.; Pike, A. C. W.; Chang, Y.-N.; Prakaash, D.; Gelova, Z.; Stanka, J.; Moreau, C.; Scott, H. C.; Wunder, F.; Wolf, G.; Scacioc, A.; McKinley, G.; Batoulis, H.; Mukhopadhyay, S.; Garofoli, A.; Pinto-Fernández, A.; Kessler, B. M.; Burgess-Brown, N. A.; Štefanić, S.; Wiedmer, T.; Dürr, K. L.; Puetter, V.; Ehrmann, A.; Khalid, S.; Ingles-Prieto, A.; Superti-Furga, G.; Sauer, D. B. Transport and inhibition of the sphingosine-1-phosphate exporter SPNS2. Nature Communications 2025, 16, 721.

[33] Honoré, E. The neuronal background K2P channels: focus on TREK1. Nature Reviews Neuroscience 2007, 8, 251–261.

[34] Enyedi, P.; Czirják, G. Molecular background of leak K+ currents: two-pore domain potassium channels. Physiological Reviews 2010, 90, 559–605.

[35] Schmidpeter, P. A.; Petroff, J. T.; Khajoueinejad, L.; Wague, A.; Frankfater, C.; Cheng, W. W.; Nimigean, C. M.; Riegelhaupt, P. M. Membrane phospholipids control gating of the mechanosensitive potassium leak channel TREK1. Nature Communications 2023, 14, 1077.

[36] Sorum, B.; Docter, T.; Panico, V.; Rietmeijer, R. A.; Brohawn, S. G. Tension activation of mechanosensitive two-pore domain K+ channels TRAAK, TREK-1, and TREK-2. Nature Communications 2024, 15, 3142.

[37] Chemin, J.; Patel, A. J.; Duprat, F.; Lauritzen, I.; Lazdunski, M.; Honoré, E. A phospholipid sensor controls mechanogating of the K+ channel TREK-1. EMBO Journal 2005, 24, 44–53.

[38] Pang, B.; Yu, L.; Li, T.; Jiao, H.; Wu, X.; Wang, J.; He, R.; Zhang, Y.; Wang, J.; Hu, H.; Dai, W.; Chen, L.; Ren, R. Molecular basis of Spns2-facilitated sphingosine-1-phosphate transport. Cell Research 2024, 34, 173–176.

[39] Anandakrishnan, R.; Aguilar, B.; Onufriev, A. V. H++ 3.0: automating pK prediction and the preparation of biomolecular structures for atomistic molecular modeling and simulation. Nucleic Acids Research 2012, 40, W537–W541.

[40] Olsson, M. H.; Søndergaard, C. R.; Rostkowski, M.; Jensen, J. H. PROPKA3: Consistent Treatment of Internal and Surface Residues in Empirical pKa Predictions. Journal of Chemical Theory and Computation 2011, 7, 525–537.

[41] Dastvan, R.; Rasouli, A.; Dehghani-Ghahnaviyeh, S.; Gies, S.; Tajkhorshid, E. Proton-driven alternating access in a spinster lipid transporter. Nature Communications 2022, 13, 5161.

[42] Riel, E. B.; Jürs, B. C. J.; Cordeiro, S.; Musinszki, M.; Schewe, M.; Baukrowitz, T. The versatile regulation of K2P channels by polyanionic lipids of the phosphoinositide and fatty acid metabolism. Journal of General Physiology 2022, 154, e202112989.

[43] Chemin, J.; Patel, A. J.; Duprat, F.; Sachs, F.; Lazdunski, M.; Honoré, E. Upand downregulation of the mechano-gated K2P channel TREK-1 by PIP2 and other membrane phospholipids. Pflügers Archiv European Journal of Physiology 2007, 455, 97–103.

[44] Cabanos, C.; Wang, M.; Han, X.; Hansen, S. B. A Soluble Fluorescent Binding Assay Reveals PIP2 Antagonism of TREK-1 Channels. Cell Reports 2017, 20, 1287–1294.

[45] Panasawatwong, A.; Pipatpolkai, T.; Tucker, S. J. Transition between conformational states of the TREK-1 K2P channel promoted by interaction with PIP2. Biophysical Journal 2022, 121, 2380–2388.

[46] Li, J.; Wang, L.; Zhu, Z.; Song, C. Exploring the alternative conformation of a known protein structure based on contact map prediction. Journal of Chemical Information and Modeling 2024, 64, 301–315.

[47] Guan, X.; Tang, Q.-Y.; Ren, W.; Chen, M.; Wang, W.; Wolynes, P. G.; Li, W. Predicting protein conformational motions using energetic frustration analysis and AlphaFold2. Proceedings of the National Academy of Sciences 2024, 121, e2410662121.

[48] Karanicolas, J.; Brooks III, C. L. Improved Gō-like models demonstrate the robustness of protein folding mechanisms towards non-native interactions. Journal of molecular biology 2003, 334, 309–325.

[49] Li, W.; Wang, W.; Takada, S. Energy landscape views for interplays among folding, binding, and allostery of calmodulin domains. Proceedings of the National Academy of Sciences 2014, 111, 10550–10555.

[50] Vickery, O. N.; Stansfeld, P. J. CG2AT2: An enhanced fragment-based approach for serial multi-scale molecular dynamics simulations. Journal of chemical theory and computation 2021, 17, 6472–6482.

[51] Rizo, J.; Sari, L.; Jaczynska, K.; Rosenmund, C.; Lin, M. M. Molecular mechanism underlying SNARE-mediated membrane fusion enlightened by all-atom molecular dynamics simulations. Proceedings of the National Academy of Sciences 2024, 121, e2321447121.

[52] Denesyuk, N. A.; Thirumalai, D. Coarse-grained model for predicting RNA folding thermodynamics. Journal of Physical Chemistry B 2013, 117, 4901–4911.

[53] Knotts 4th, T. A.; Rathore, N.; Schwartz, D. C.; de Pablo, J. J. A coarse grain model for DNA. Journal of Chemical Physics 2007, 126, 084901.

[54] Gil Herrero, C.; Duve, T.; Thallmair, S. Martini 3 coarse-grained models of azobenzene-based photolipids: modulation of membranes with light. Journal of Chemical Theory and Computation 2026, 22, 708–722.

[55] MacCallum, J. L.; Hu, S.; Lenz, S.; Souza, P. C.; Corradi, V.; Tieleman, D. P. An implementation of the Martini coarse-grained force field in OpenMM. Biophysical Journal 2023, 122, 2864–2870.

[56] Izaguirre, J. A.; Sweet, C. R.; Pande, V. S. Multiscale dynamics of macromolecules using normal mode langevin. Pacific Symposium on Biocomputing 2010, 240–251.

[57] Chow, K. H.; Ferguson, D. M. Isothermal-isobaric molecular dynamics simulations with Monte Carlo volume sampling. Computer Physics Communications 1995, 91, 283–289.

[58] Åqvist, J.; Wennerström, P.; Nervall, M.; Bjelic, S.; Brandsdal, B. O. Molecular dynamics simulations of water and biomolecules with a Monte Carlo constant pressure algorithm. Chemical Physics Letters 2004, 384, 288–294.

[59] Bussi, G.; Donadio, D.; Parrinello, M. Canonical sampling through velocity rescaling. The Journal of chemical physics 2007, 126.

[60] Parrinello, M.; Rahman, A. Polymorphic transitions in single crystals: A new molecular dynamics method. Journal of Applied physics 1981, 52, 7182–7190.

[61] Tironi, I. G.; Sperb, R.; Smith, P. E.; Gunsteren, W. F. V. A generalized reaction field method for molecular dynamics simulations. The Journal of Chemical Physics 1995, 102, 5451–5459.

[62] Wołek, K.; Gómez-Sicilia, À.; Cieplak, M. Determination of contact maps in proteins: A combination of structural and chemical approaches. The Journal of Chemical Physics 2015, 143, 243105.

[63] Gronenborn, A. M.; Filpula, D. R.; Essig, N. Z.; Achari, A.; Whitlow, M.; Wingfield, P. T.; Clore, G. M. A Novel, Highly Stable Fold of the Immunoglobulin Binding Domain of Streptococcal Protein G. Science 1991, 253, 657–661.

[64] Vijay-Kumar, S.; Bugg, C. E.; Cook, W. J. Structure of ubiquitin refined at 1.8 Å resolution. Journal of Molecular Biology 1987, 194, 531–544.

[65] Wassenaar, T. A.; Ingólfsson, H. I.; Böckmann, R. A.; Tieleman, D. P.; Marrink, S. J. Computational lipidomics with insane: A versatile tool for generating custom membranes for molecular simulations. Journal of Chemical Theory and Computation 2015, 11, 2144–2155.

[66] Bjorkman, A.; Mowbray, S. Multiple open forms of ribose-binding protein trace the path of its conformational change. Journal of Molecular Biology 1998, 279, 651–664.

[67] Bjorkman, A.; Binnie, R.; Zhang, H.; Cole, L.; Hermodson, M.; Mowbray, S. Probing proteinprotein interactions. The ribose-binding protein in bacterial transport and chemotaxis. Journal of Biological Chemistry 1994, 269, 30206–30211.

[68] Šali, A.; Blundell, T. L. Comparative protein modelling by satisfaction of spatial restraints. Journal of Molecular Biology 1993, 234, 779–815.

[69] Pedersen, K. B.; Ingólfsson, H. I.; Ramirez-Echemendia, D. P.; Borges-Araújo, L.; Andreasen, M. D.; Empereur-Mot, C.; Melcr, J.; Ozturk, T. N.; Bennett, W. F. D.; Kjølbye, L. R.; Brasnett, C.; Corradi, V.; Khan, H. M.; Cino, E. A.; Crowley, J.; Kim, H.; Fábián, B.; Borges-Araújo, A. C.; Pavan, G. M.; Launay, G.; Lolicato, F.; Wassenaar, T. A.; Melo, M. N.; Thallmair, S.; Carpenter, T. S.; Monticelli, L.; Tieleman, D. P.; Schiøtt, B.; Souza, P. C. T.; Marrink, S. J. The Martini 3 Lipidome: Expanded and Refined Parameters Improve Lipid Phase Behavior. ACS Central Science 2025, 11, 1598–1610.

[70] Michaud-Agrawal, N.; Denning, E. J.; Woolf, T. B.; Beckstein, O. MDAnalysis: A toolkit for the analysis of molecular dynamics simulations. Journal of Computational Chemistry 2011, 32, 2319–2327.

[71] Pettersen, E. F.; Goddard, T. D.; Huang, C. C.; Meng, E. C.; Couch, G. S.; Croll, T. I.; Morris, J. H.; Ferrin, T. E. UCSF ChimeraX: Structure visualization for researchers, educators, and developers. Protein Science 2021, 30, 70–82.

