## Supplemental File for "*CTGoMartini*: A Python Framework for Simulating Biomolecular Conformational Transitions with Gō-Martini Models"

### 1 Supplementary Tables

**Table S1:** Comparison results of single-point energy and force calculations between GROMACS and CTGoMartini.

| Name | Description | Energy | | Force | |
|---|---|---|---|---|---|
| | | Absolute error (kJ/mol) | Relative error | Absolute error (kJ/mol/nm) | Relative error |
| KALP | Antimicrobial peptide | 5.1e-06 | 2.6e-11 | 4.1e-07 | 6.0e-07 |
| 1GB1 | Domain of protein G | 6.5e-06 | 1.8e-11 | 3.8e-07 | 1.6e-07 |
| 1UBQ | Ubiquitin | 6.8e-06 | 1.9e-11 | 4.0e-06 | 1.9e-06 |
| 1GGG | Open state of GlnBP | 2.8e-06 | 2.6e-11 | 3.4e-06 | 1.3e-06 |
| 1WDN | Closed state of GlnBP | 2.8e-06 | 2.6e-11 | 2.5e-06 | 5.2e-07 |
| 1ANK | Closed state of AdK | 3.9e-06 | 2.8e-11 | 3.5e-06 | 6.2e-07 |
| 3SN6 | Active state of Beta2AR | 3.5e-06 | 2.9e-11 | 5.7e-06 | 9.5e-07 |
| 8DE9 | Down state of TREK1 | 8.5e-06 | 3.2e-11 | 3.2e-06 | 1.6e-06 |

#### 2 Supplementary Figures

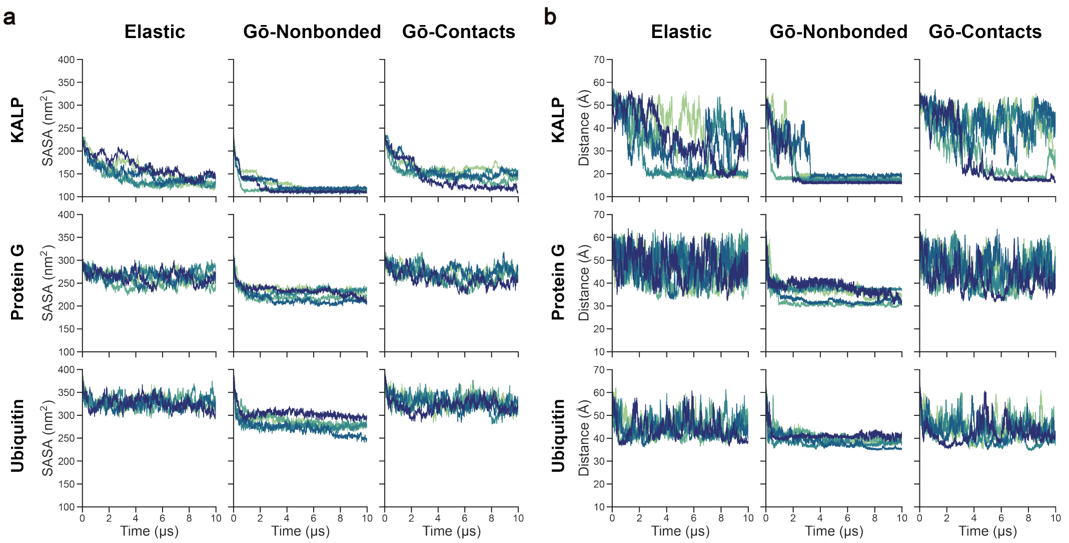


**Figure S1: Time evolution of protein aggregations.** Solvent accessible surface area (SASA) (**a**) and averaged intermolecular distance (**b**) plotted as a function of simulation time. The rows correspond to KALP, Protein G, and Ubiquitin systems, while the columns display results for the Elastic, Gō-Nonbonded, and Gō-Contacts models. Each plot contains trajectories from five independent replicates, shown in distinct colors.

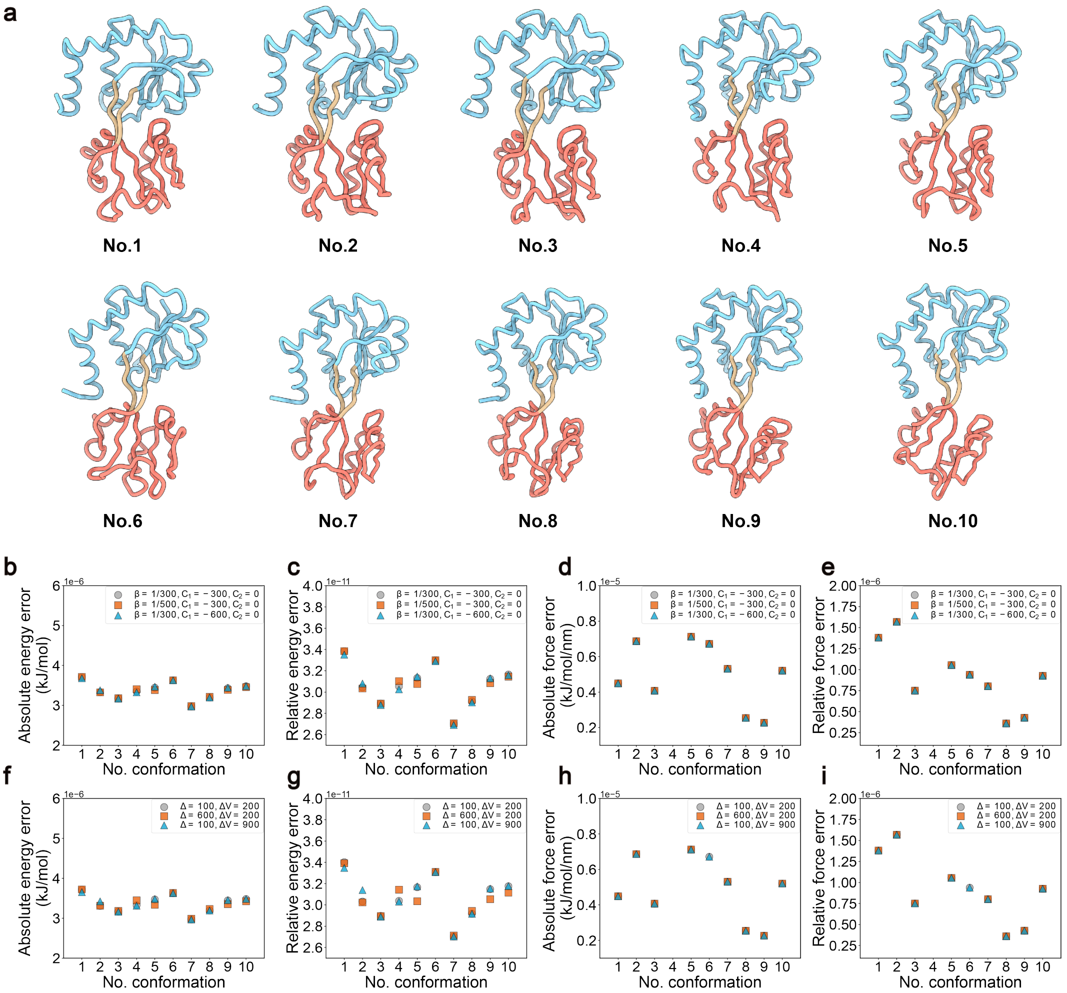


**Figure S2: Validation of energy and force calculations for Multiple-basin Gō-Martini using the GlnBP system. a,** Ten closed-to-open representative conformations of GlnBP used for the test. **b-e,** Absolute and relative errors for the calculated energy and forces under different EXP parameters. **f-i,** Absolute and relative errors for the calculated energy and forces under different HAM parameters.

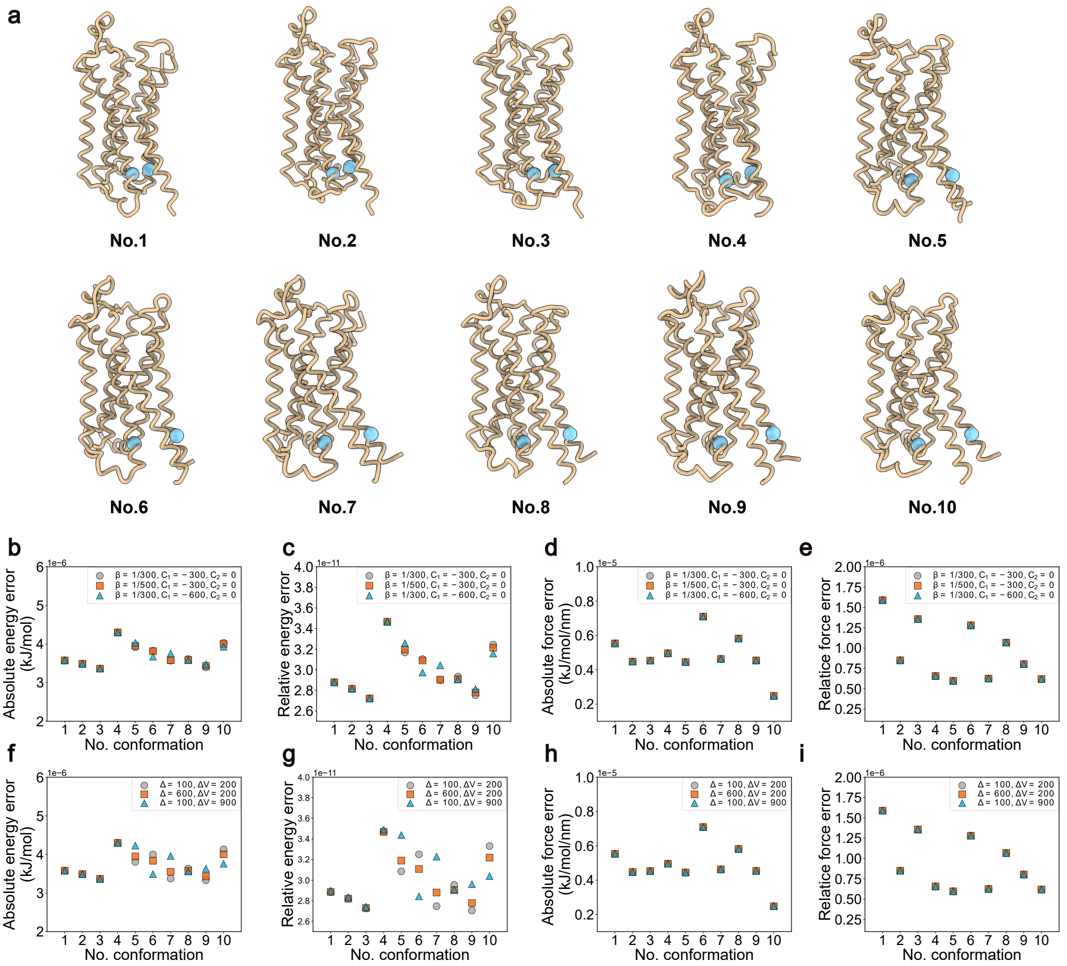


**Figure S3: Validation of energy and force calculations for Multiple-basin Gō-Martini using the $\beta_2$AR system.** **a,** Ten inactive-to-active representative conformations of $\beta_2$AR used for the test. BB of R131 and L272 are colored sky-blue with sphere style. **b-e,** Absolute and relative errors for the calculated energy and forces under different EXP parameters. **f-i,** Absolute and relative errors for the calculated energy and forces under different HAM parameters.

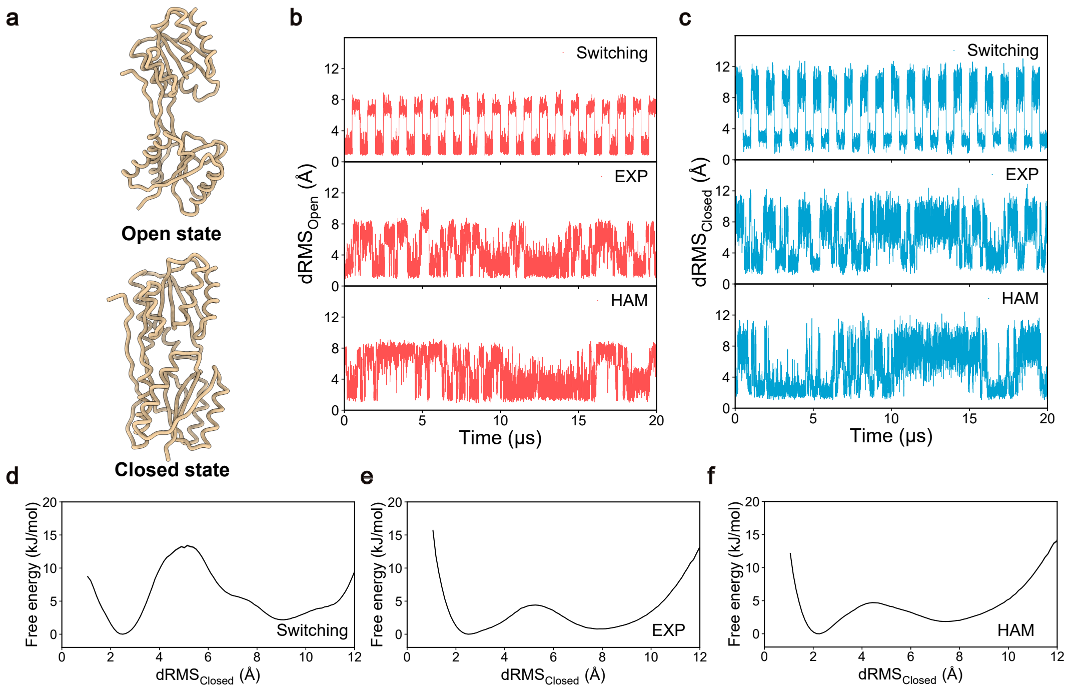


**Figure S4: Comparison of Switching and Multiple-basin Gō-Martini approaches for simulating RBP conformational transitions. a,** Coarse-grained structures of RBP in the open (PDB ID: 1BA2) and closed (PDB ID: 2DRI) states. **b, c,** Time evolution of dRMS with respect to the open state (**b**) and closed state (**c**) over 20 μs simulations. **d-f,** Free energy profiles of RBP as a function of $dRMS_{Closed}$ for the Switching (**d**), EXP (**e**), and HAM (**f**) methods, respectively. Profiles are derived from 10 independent replicates, with the first 1 μs of each trajectory discarded during the analysis. Shaded areas represent the statistical uncertainty estimated using bootstrap analysis. Note that these errors are negligibly small and thus not visible in the figure.

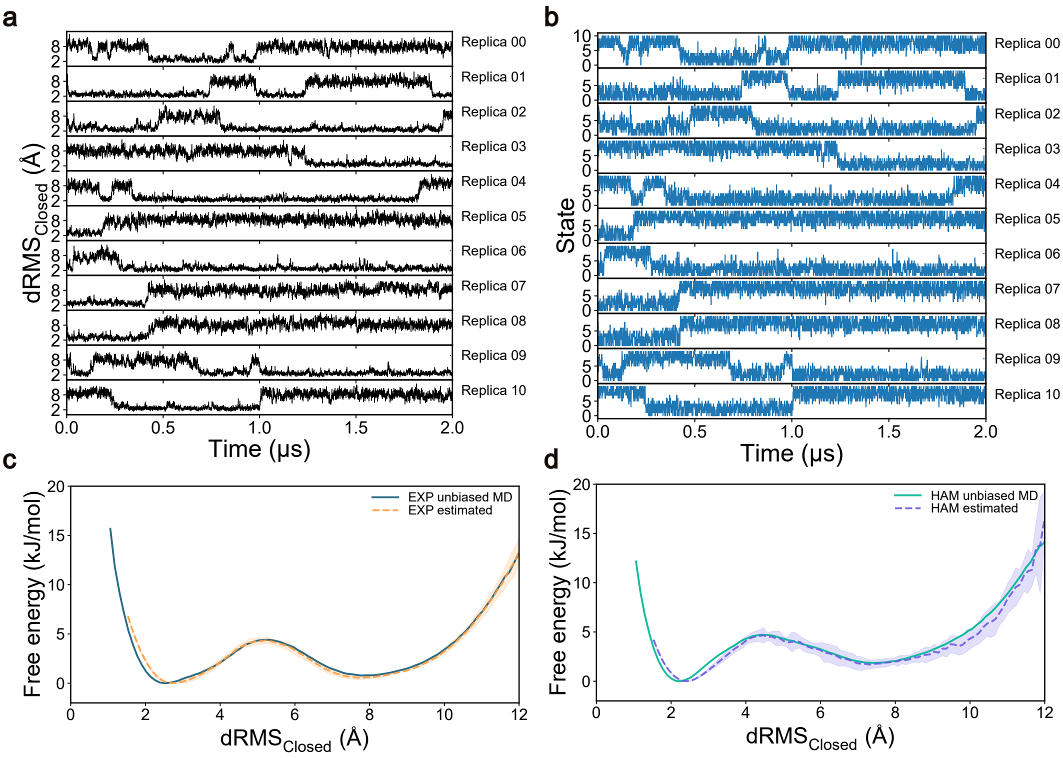


**Figure S5: Parameter optimization of Multiple-basin Gō-Martini models of RBP. a,** Time evolution of the $dRMS_{Closed}$ of RBP for 11 individual replicas over a 2 μs simulation. The fluctuations indicate transitions between distinct conformational states. **b,** Time evolution of the Hamiltonian state index of RBP for each replica. The rapid variation in state indices demonstrates efficient mixing and diffusion of replicas across the Hamiltonian ladder. **c, d,** Comparison of free energy profiles derived from HREMD simulations via PyMBAR predictions (dashed lines) versus long unbiased MD simulations (solid lines). Results are shown for the EXP mixing method (**c**) and the HAM mixing method (**d**). All predictions are derived from PyMBAR using raw HREMD data collected using the EXP mixing method. The shaded regions represent statistical uncertainty calculated from three independent replicates.

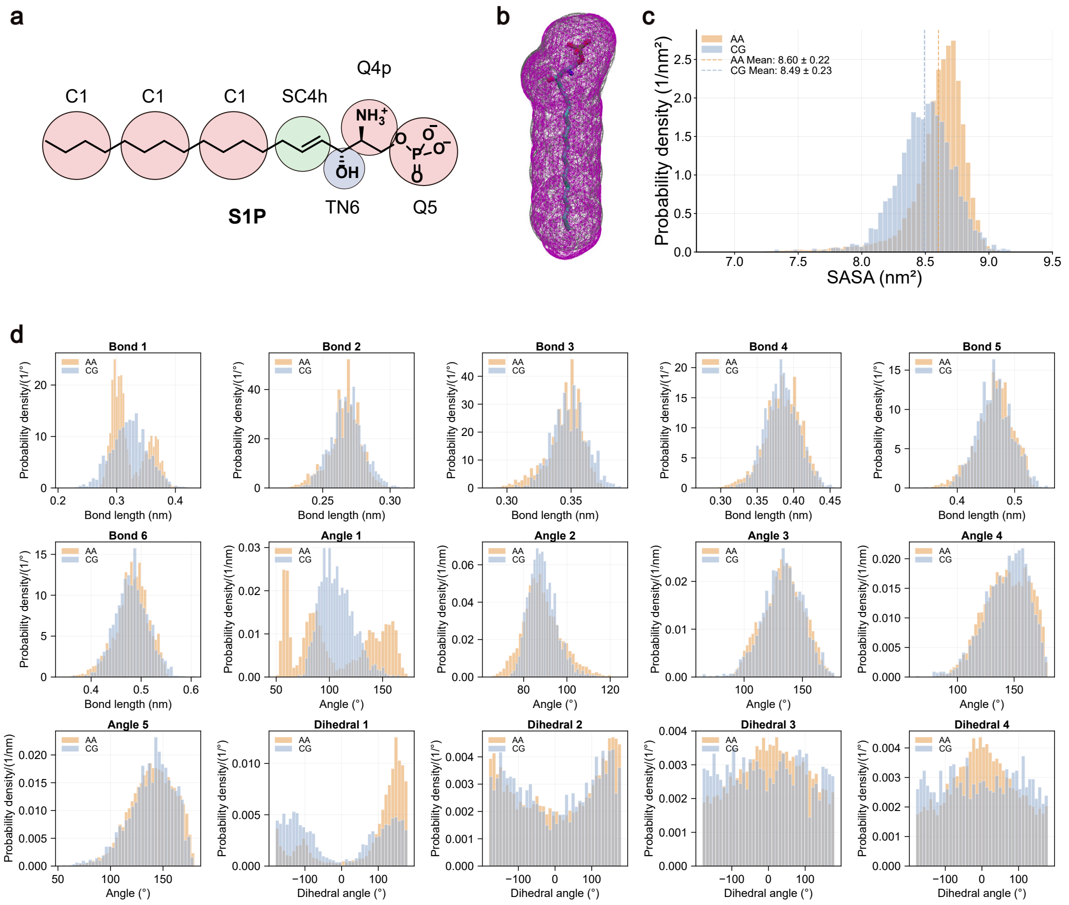


**Figure S6: Parameterization and validation of the coarse-grained S1P model. a,** Schematic illustration of the mapping from the atomistic structure to the coarse-grained representation. **b,** Volumetric comparison showing the van der Waals surface superposition of the all-atom and coarse-grained models. **c,** Probability density distributions of SASA for both the all-atom and coarse-grained models. **d,** Probability densities for bond lengths, angles, and dihedrals are plotted for both the all-atom and coarse-grained trajectories.

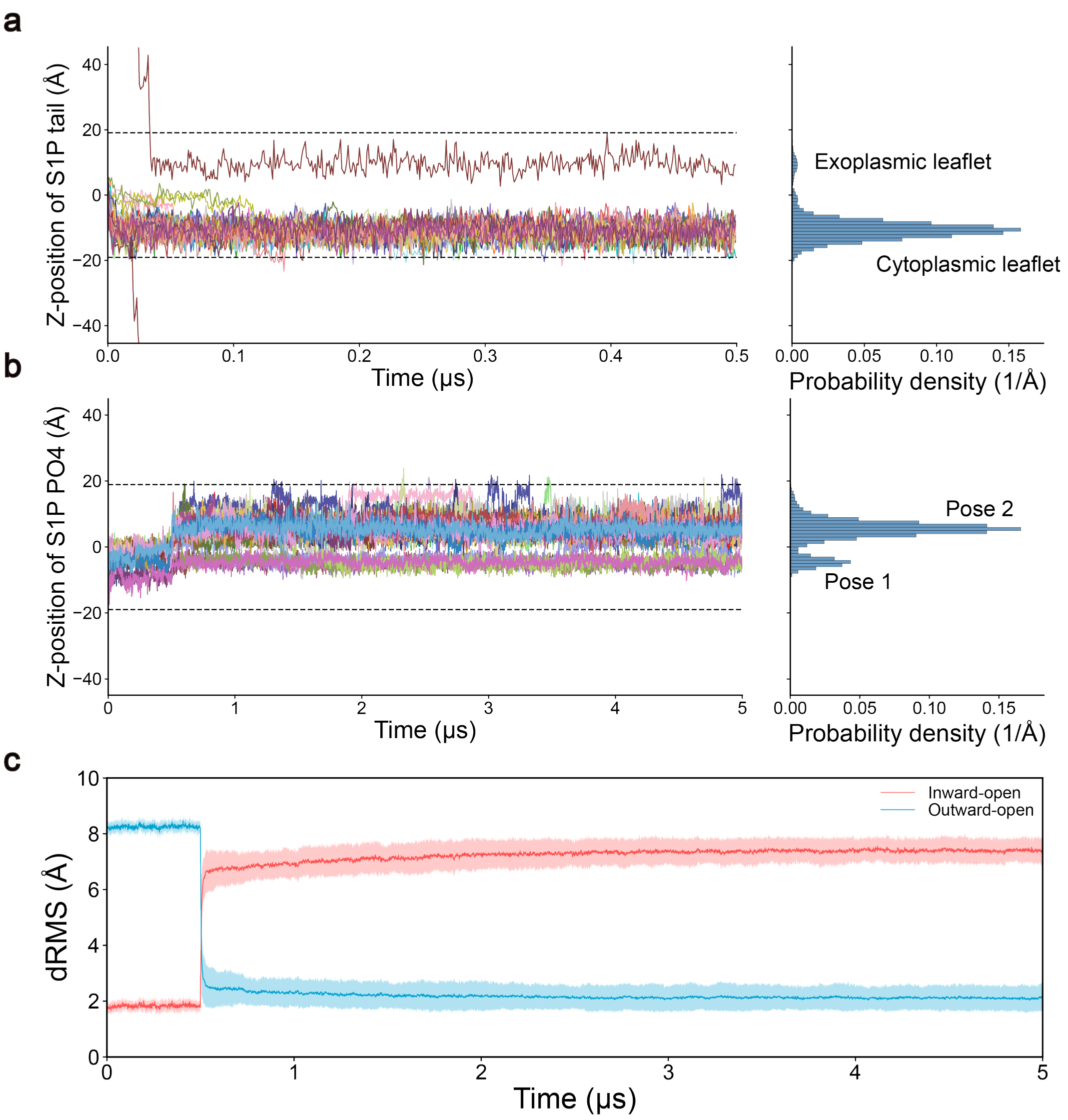


**Figure S7: Dynamics of S1P induced by SPNS2 conformational transition. a,** Time evolution and probability density distribution of the Z-position of the S1P hydrophobic tail for the unbinding events. **b,** Time evolution and probability density distribution of the Z-position of the S1P phosphate head group. The data are from the trajectories of S1P binding but not translocated events. **c,** Time evolution of dRMS relative to the Inward-open (red) and Outward-open (blue) reference structures. The conformational transitions from the Inward-open to the Outward-open state induced by the Switching method are observed at approximately 0.5 µs. Shaded regions indicate the standard deviation from 52 independent replicates.

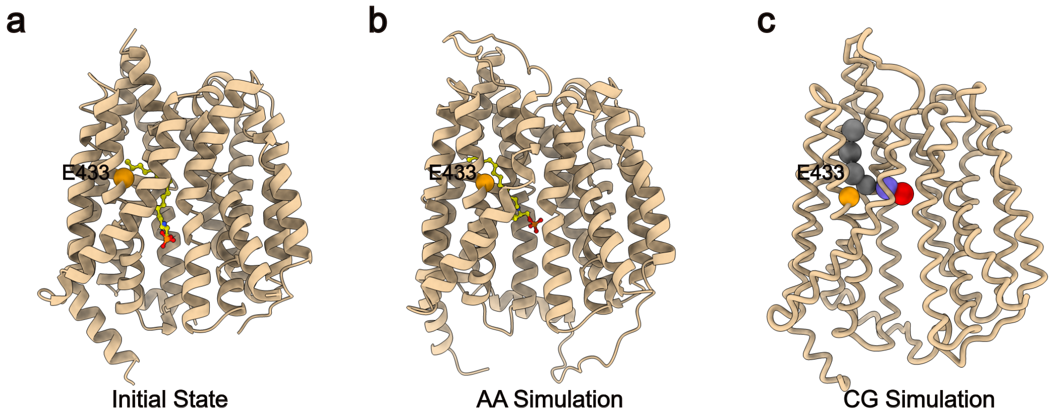


**Figure S8: Comparison of S1P binding poses in the inward-open state of SPNS2. a,** Initial binding pose from the PDB structure (PDB ID: 8EX4). **b,** Binding pose from atomistic molecular dynamics simulations.[1] **c,** Binding pose from CG simulations with CTGoMartini. In all panels, S1P is shown as a ball-and-stick model, and the CA/BB bead of E433 is highlighted as an orange sphere.

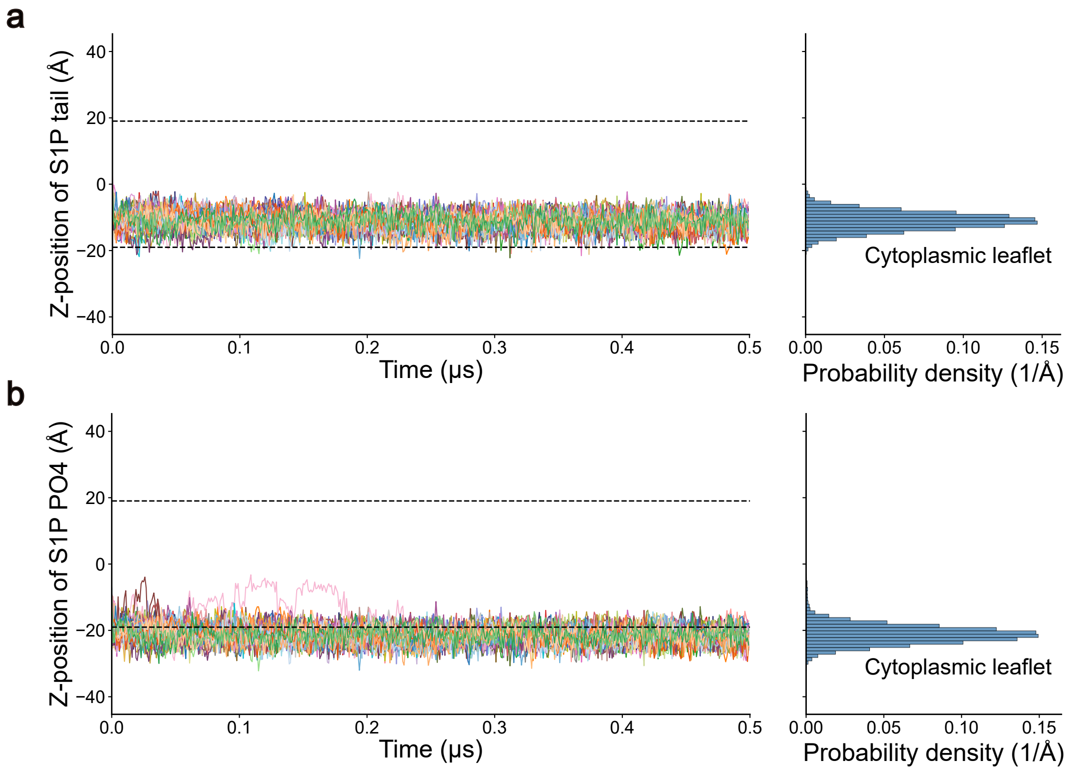


**Figure S9: Dynamics of S1P transport in SPNS2 with deprotonated E433. a,** Time evolution (left) and probability density distribution (right) of the Z-position of the S1P tail. **b,** Time evolution (left) and probability density distribution (right) of the Z-position of the S1P phosphate group.

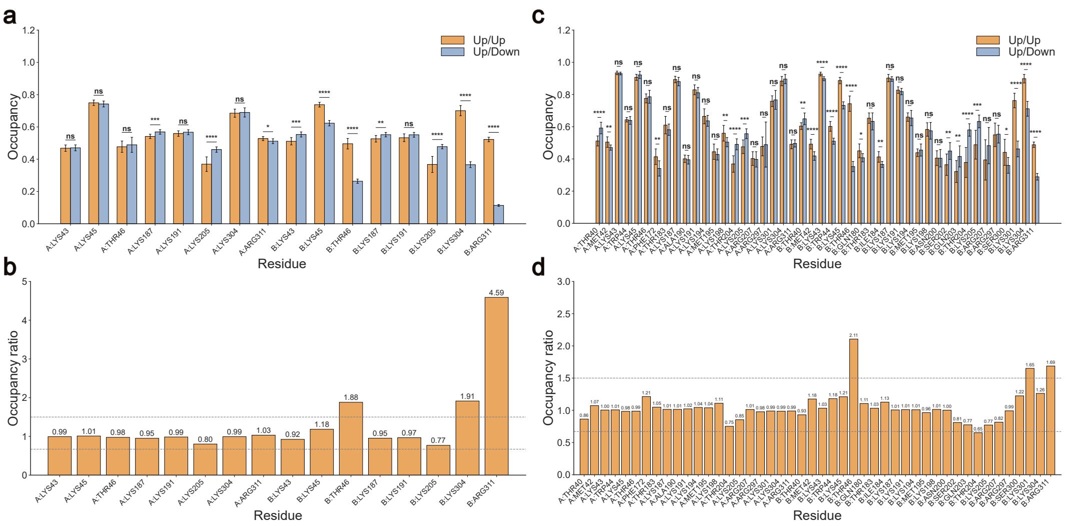


**Figure S10: Quantitative analysis of state-dependent lipid-protein interactions. a,** POPA lipid occupancy for key residues in the Up/Up and Up/Down states. Only residues exhibiting a lipid occupancy greater than 40% in at least one of the two states are displayed. Statistical significance is determined using a two-tailed t-test (ns: not significant, $*p < 0.05$, $**p < 0.01$, $***p < 0.001$, $****p < 0.0001$). **b,** The ratio of lipid occupancy in the Up/Up state relative to the Up/Down state for the residues shown in **a**. Dashed lines indicate ratio thresholds of 0.67 and 1.5. **c,** PIP2 lipid occupancy for key residues in the Up/Up and Up/Down states. Selection criteria and statistical analysis are identical to panel **a**. **d,** The ratio of lipid occupancy in the Up/Up state relative to the Up/Down state for the residues shown in **c**. Dashed lines indicate ratio thresholds of 0.67 and 1.5.

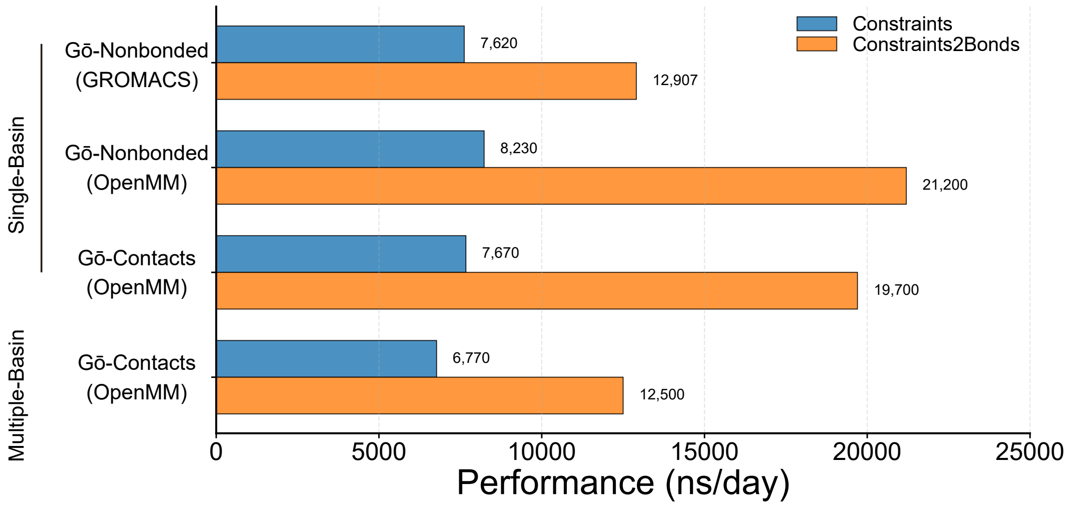


**Figure S11: Performance comparison of CTGoMartini (OpenMM) and GROMACS for different Gō-Martini models.** Simulation speeds are compared using the GlnBP system with different implementations. Comparisons include: (i) the single-basin Gō-Nonbonded model in GROMACS versus OpenMM with both standard constraints and constraints converted to bonds, (ii) the single-basin Gō-Contacts model in OpenMM with both constraint treatments, and (iii) the multiple-basin Gō-Martini model (EXP mixing scheme) with standard constraints versus with constraints converted to bonds. For the bond conversion, a harmonic force constant of 50,000 kJ/mol·nm² was used. All benchmarks were performed on a workstation with an NVIDIA GeForce RTX 4080 GPU and an Intel Core i7-14700K CPU.

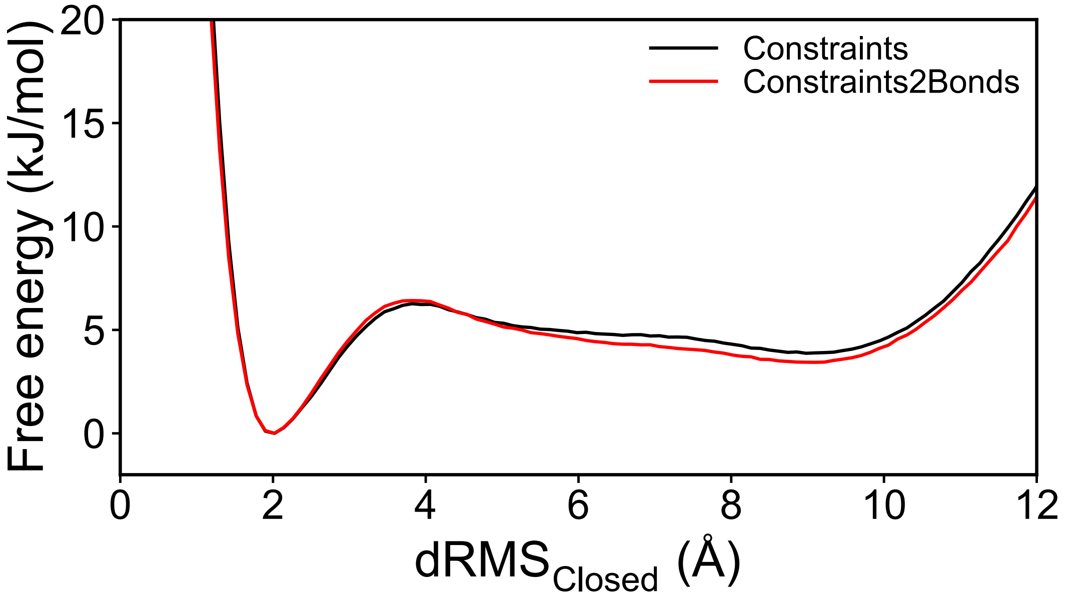


**Figure S12: Comparison of free energy profiles for GlnBP with standard constraints versus constraints converted to bonds.** Free energy as a function of $\mathrm{dRMS}_{\mathrm{Closed}}$ for simulations using standard constraints (black) and with constraints converted to bonds (red). Each profile was computed from 10 independent 20-μs simulations, with the first 1 μs of each trajectory discarded during the analysis. For the bond conversion, a harmonic force constant of 50,000 kJ/mol·nm$^2$ was used.

### 3 Supplementary Movie

**Supplementary Movie 1. Translocation process of S1P mediated by the conformational transitions of SPNS2**. The movie presents a representative MD trajectory illustrating the S1P translocation process, which is coupled to the conformational switching of SPNS2 from the inward-open to outward-open state. The SPNS2 protein is rendered in tan, and S1P is depicted as a ball-and-stick model.

### References

[1] Li, H. Z. et al. Transport and inhibition of the sphingosine-1-phosphate exporter SPNS2. *Nature Communications* **2025**, *16*, 721.
